# A mouse model of autosomal dominant spastic ataxia and myopathy caused by a mutation in *Tuba4a*

**DOI:** 10.64898/2026.03.06.710113

**Authors:** Timothy J. Hines, Jonathan R. Funke, Samia L. Pratt, Alaura D. Rice, Jeffery L. Twiss, Robert W. Burgess

## Abstract

Hereditary ataxias are a heterogeneous group of neurodegenerative disorders characterized by impaired balance and coordination, often due to cerebellar dysfunction. Despite advances in identifying genetic causes, animal models remain essential for dissecting underlying mechanisms and testing therapeutic strategies. Here we describe a mouse model of spastic ataxia and myopathy caused by a missense mutation in *Tuba4a* (n.A626C, p.Gln176Pro). In an ENU mutagenesis screen, a male C57BL/6J mouse exhibiting muscle wasting and an intention tremor starting at approximately 4 weeks-of-age was identified. The male was bred by in vitro fertilization to BALB/cByJ oocyte donors. Genetic mapping determined dominant inheritance and localized the mutation to Chromosome 1. Genome sequencing revealed single nucleotide polymorphisms (SNPs) in serine threonine kinase 36 (*Stk36^Y1003N^*) and alpha-tubulin 4A (*Tuba4a^Q176P^*) in the mapping interval. These SNPs were CRISPR-engineered into C57BL/6J mice, which confirmed the *Tuba4a^Q176P^* variant as the causative mutation. Mutant mice are normal at 3 weeks, except for decrement in muscle response following repetitive nerve stimulation. However, by 30 days these mice have ataxia, Purkinje neuron degeneration, and extensive skeletal muscle defects, which contribute to a decreased lifespan. Dominant *TUBA4A* mutations in humans are associated with spastic ataxia type 11 (SPAX11), congenital myopathy type 26 (CMYO26), and frontotemporal dementia/amyotrophic lateral sclerosis type 9 (FTDALS9). Our mice exhibit hallmark features of SPAX11 and CMYO26, but do not show motor neuron degeneration. This specificity makes this model a valuable tool for studying cell-type selective effects of *TUBA4A* mutations in neurodegeneration and myopathy.

## INTRODUCTION

Microtubules are part of the cytoskeleton, a dynamic meshwork of filamentous protein complexes that provide structural support and spatial organization within cells. In addition to their structural role, microtubules are critical for cell migration during development, form the mitotic spindle during cell division, contribute to cell motility as major components of cilia and flagella, and serve as “tracks” for intracellular transport mediated by motor proteins such as kinesins and dyneins^1^. Microtubules consist of linear chains of polymerized α-/β-tubulin heterodimers called protofilaments, which bundle together to form the hollow tube structure of the microtubule^1,2^. Although they are primarily composed of only two proteins, microtubules can be highly specialized due to a wide range of microtubule-associated proteins, as well as multiple genetically distinct subtypes and posttranslational modifications for both α- and β-tubulin, which alter microtubule stability, dynamics, and function^1,2^.

The human genome contains 10 α-tubulin (*TUBA*s) and 9 β-tubulin (*TUBB*s) genes, while mice have 8 of each^3^. Of these, six α-tubulin and seven β-tubulin genes are shared between the two species (Table 1). Tubulin genes can have specialized functions and tissue-specific expression patterns, adding complexity to their biology. Mutations in tubulin genes cause a group of disorders collectively known as tubulinopathies, which can present with a wide spectrum of clinical features affecting different organ systems^4,5^. Despite this variability, the nervous system is often affected by these conditions. For example, mutations in *TUBA1A*, *TUBA8*, *TUBB2A*, *TUBB2B*, *TUBB3*, and *TUBB5* lead to neurodevelopmental conditions such as polymicrogyria, lissencephaly, and other brain malformations, while mutations in *TUBA4A*, *TUBB3*, and *TUBB4A* are associated with neurodegenerative disorders including frontotemporal dementia with amyotrophic lateral sclerosis (FTDALS), spastic ataxia, axonal neuropathy, and hypomyelination with atrophy of basal ganglia and cerebellum (H-ABC)^4–17^.

**Table 1.**
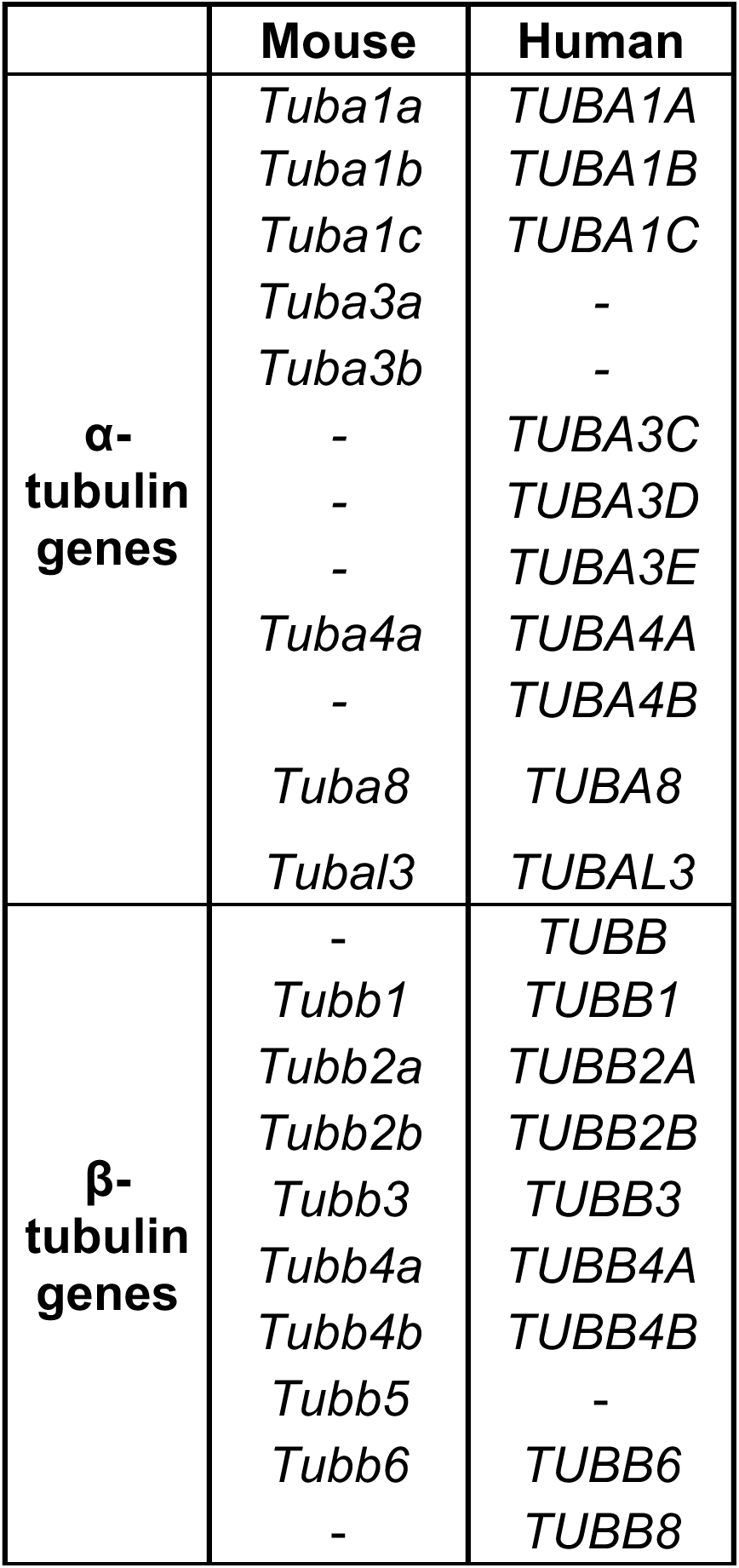
Mouse and human α- and β-tubulin genes.

In some cases, different mutations in the same tubulin gene can cause dramatically different phenotypes. For example, dominant mutations in *TUBA4A* are associated with Congenital Myopathy type 26 (CMYO26), spastic ataxia type 11 (SPAX11), frontotemporal dementia with or without ALS type 9 (FTDALS9), and potentially Parkinson’s disease (PD), disorders which primarily affect skeletal muscle, the cerebellum, cortical and spinal motor neurons, and dopaminergic neurons in the substantia nigra, respectively ^9–12,16–20^. However, there is clinical and genetic overlap between these neurodegenerative disorders, including SPAX11 patients with cognitive dysfunction, FTD patients with parkinsonism, and a common mutation (*TUBA4A^R105C^*) associated with both SPAX11 and FTDALS9^9,16^. In addition to neurological and muscular disorders, dominant and recessive *TUBA4A* mutations are also associated with oocyte, zygote, embryo maturation arrest type 23 (OZEMA23) and macrothrombocytopenia^21–23^.

In this paper, we describe a mouse strain isolated from an ENU mutagenesis screen that presents with a “jerky” gait and skeletal muscle pathology around 30 days. This gait phenotype led to the original strain nickname “Jerky”. However, that name is already in use for an epileptic mouse strain^24^, so the strain described in this manuscript has been given the related name, “Biltong”. Genetic mapping and genome sequencing revealed candidate single nucleotide polymorphisms (SNPs) in the coding sequences of serine threonine kinase 36 (*Stk36^Y1003N^*) and alpha-tubulin 4A (*Tuba4a^Q176P^*). CRISPR-engineering these variants into C57BL/6J mice confirmed *Tuba4a^Q176P^* as the causative mutation. This mutation is located three amino acids away from a CMYO26-associated variant (T179I) and two SPAX11-associated variants (P173R and P173S)^16,18^. The mice described here have phenotypes consistent with both diseases, including Purkinje cell degeneration in the cerebellum leading to ataxia, and skeletal muscle pathology such as myofibrillar disorganization, inclusions, and vacuoles. Notably, the mice lack ALS-associated pathology such as loss of spinal motor neurons or their axons, making this a valuable model for studying cell type specific effects of pathogenic *TUBA4A* mutations.

## RESULTS

### Identification of the *Biltong* mutation

A 4-week old C57BL/6J (B6) male from an ENU mutagenesis screen presented with jerky movements, decreased locomotion, and muscle atrophy. Attempts at conventional breeding were unsuccessful; therefore, the line was propagated via in vitro fertilization using wild-type BALB/cByJ (BALB) oocyte donors. Roughly half of the F1 offspring inherited the phenotype, indicating a dominant mode of inheritance. Affected offspring were backcrossed to BALB to generate N2 mice for genetic mapping. The line was maintained on a mixed B6/BALB background for breeding purposes (CByJ;B6J-Tuba4a<BILT>/Rwb). These mice are referred to as *Biltong* due to their “jerky” gait (Video 1 and Video 2). All genomic coordinates in this manuscript refer to build GRCm38/mm10.

To map the mutation, a genome-wide B6/BALB SNP panel with 141 markers was used to look for association of the phenotype with a SNP marker in 22 affected N2 mice and 22 unaffected littermates. One SNP on Chromosome 1 segregated completely with the phenotype: rs3677697 at 80,181,487 bp (not shown). Analysis of additional B6/BALB SNPs revealed two more segregating SNPs: rs3672814 and rs4222476 at 78,030,398 bp and 79,289,860 bp, respectively (Figure 1A). The two markers flanking these co-segregating SNPs that showed recombinations excluding more distal and proximal regions of Chr. 1 are located at 69,020,216 bp (rs3683684) and 94,081,544 bp (rs3664528), giving a ∼25 Mbp region of interest (Figure 1A). A list of genes in this interval can be found in Supplemental Table 1.

**Figure 1.**
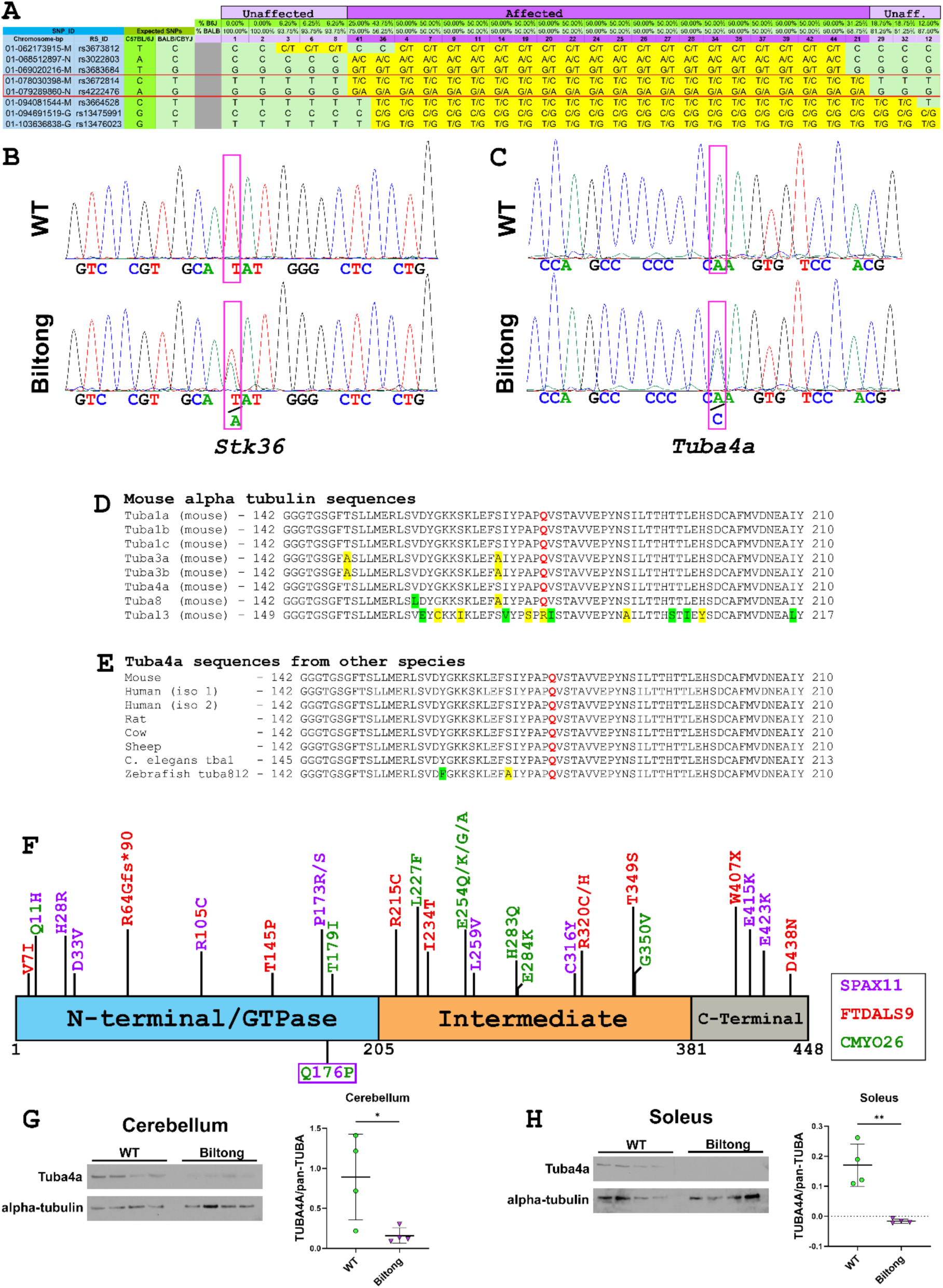
Genetic mapping and sequencing of Biltong mice. **A)** Results from fine genetic mapping experiment showing the two segregating SNPs (red box) from the panel. Sample numbers shaded with light purple are unaffected mice and those shaded with dark purple are affected. **B&C)** Sanger sequencing chromatograms showing heterozygous SNPs found in serine threonine kinase 36 (*Stk36*) and alpha tubulin 4A (*Tuba4a*) by whole exome and whole genome sequencing, respectively. **D&E)** The affected glutamic acid (red Q) is highly conserved both between different mouse alpha tubulin genes and between *Tuba4a* genes/orthologs from different species, including rat, cow, sheep, *C. elegans* (*tba1*), and zebrafish (*tuba8l2*). Residues highlighted in yellow are non-conservative substitutions and those in green are conservative substitutions. **F)** Schematic of TUBA4A protein showing dominant mutations associated with SPAX11, FTDALS9, and CMYO26 in purple, red, and green, respectively^8–12,16–19^. The R105C mutation has alternating purple and red characters because it has been associated with both SPAX11 and FTDALS9. The patient with the Q11H mutation was diagnosed with multisystem proteinopathy, but presented with ataxia and myofibrillar myopathy, so this mutation has alternating green and purple characters. The mutation identified in this study, Q176P, is in a box on the bottom with alternating green and purple characters. **G&H**) Western blots for TUBA4A and pan-alpha tubulin from Cerebellum (G) and soleus (H) of WT and Biltong mice, with quantification of band intensity.

Whole exome sequencing (WES) and whole genome sequencing (WGS) revealed candidate SNPs in the coding sequences of serine threonine kinase 36 (*Stk36^Y1003N^*) and alpha-tubulin 4a (*Tuba4a^Q176P^*) within the region of interest on Chr. 1 (Figure 1B, C). Since ENU mutagenesis typically induces A>G point mutations, which are more likely to produce a phenotype when occurring in coding sequence than in intronic or intergenic regions, we prioritized validation of these mutations. Both variants were CRISPR-engineered into B6 mice. The *Tuba4a^Q176P^* mutation reproduced the phenotypes seen in *Biltong* mice, whereas *Stk36^Y1003N^*mice had no discernible phenotype (see below: Figure 7, and Supplementary Figure 4). The *Tuba4a* glutamine residue that is altered (Q176) is highly conserved amongst mouse alpha tubulin genes (except *Tubal3*) and between *Tuba4a* genes (or orthologs) of different species even as distant as *C. elegans* (Figure 1D, E). Furthermore, Q176 is located three amino acids upstream of a CMYO26-associated variant (T179I) and three amino acids downstream of two SPAX11-associated variants (P173R and P173S) (Figure 1F).

A Chi-square analysis on all litters (N = 11 litters) carrying the ENU mutation that were assessed at 22 days and 30 days showed that affected mice appear at the expected Mendelian ratio (43 affected mice of 91, Χ^2^ = 0.275, df = 1, p = 0.6002) consistent with a dominantly inherited mutation with full penetrance. Additionally, affected males were mated with affected females in an attempt to generate homozygotes, but out of 23 offspring from 4 litters, no homozygotes were produced, suggesting embryonic lethality when the mutation is made homozygous.

Western blot analysis revealed significant decreases in TUBA4A protein levels in two affected tissues (cerebellum and soleus muscle, see Figures 4-6) from Biltong mice when normalized to total alpha tubulin levels, detected using a pan-alpha tubulin antibody (Figure 1G-H). Notably, TUBA4A was not detected in soleus of Biltong mice even when signals in WT samples were saturated.

### Neuromuscular phenotyping

Given the apparent muscle atrophy and the association of mutations in *TUBA4A* with ALS, neuromuscular phenotyping was performed at 22 days, 30 days, 9 weeks, and 6 months to track disease progression. The latest timepoint was 6 months because mice often required euthanasia at 6 – 8 months due to low body condition score and welfare concerns. A formal survival study was not performed. Neuromuscular phenotyping included body weight, muscle weight (triceps surae), muscle weight to body weight ratio (an indicator of muscle atrophy), an inverted wire hang test of grip strength and endurance, and electromyography (EMG) to measure compound muscle action potential (CMAP) amplitudes, sciatic motor nerve conduction velocity (NCV), and possible decrement in neuromuscular transmission with repetitive nerve stimulation (RNS). Body weights are reported for each sex (Figure 2A); however, inverted wire hang tests and EMG experiments were conducted on a randomly selected subset of mice from the cohorts shown in Figure 2A and sexes were pooled because we have not previously observed sex differences in these outcomes.

**Figure 2.**
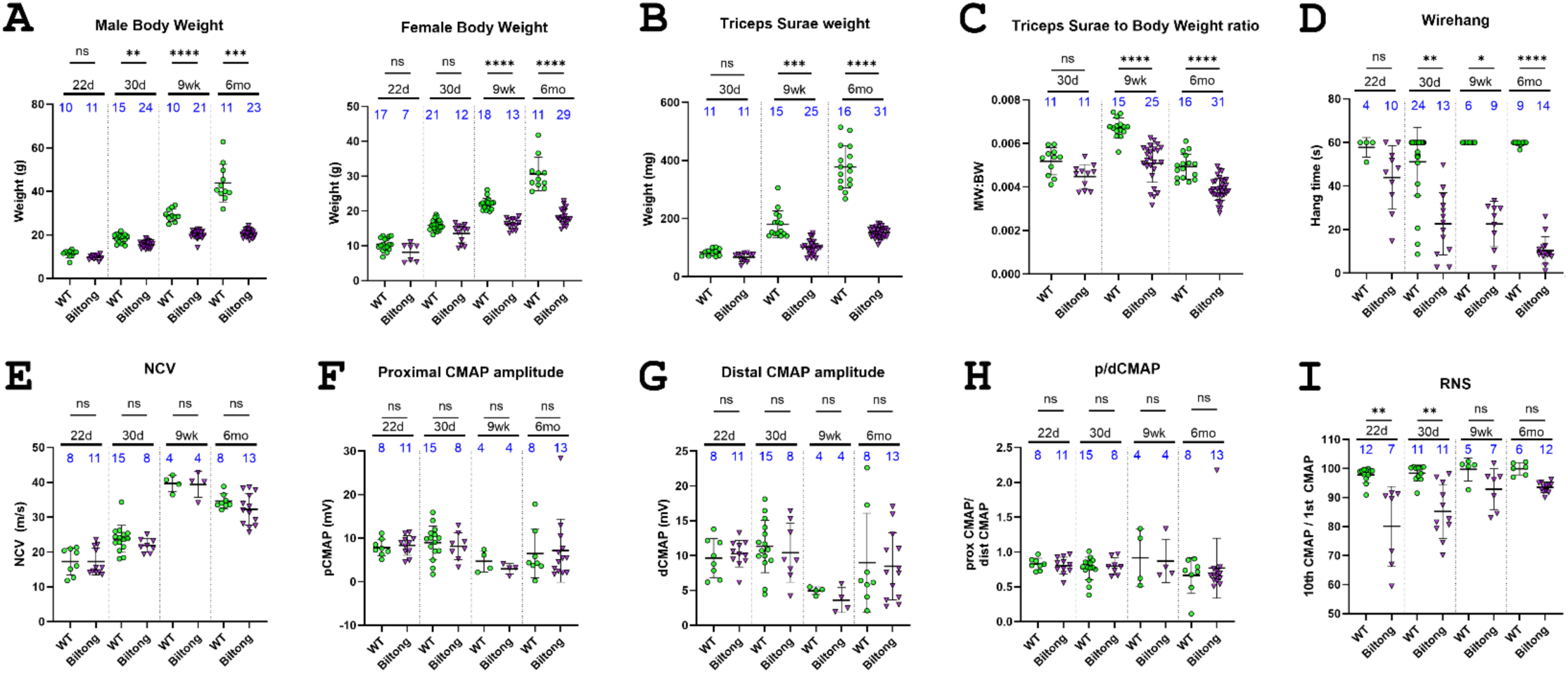
Neuromuscular phenotyping. **A)** Male and female body weights from four timepoints (22d, 30d, 9wk, and 6mo. **B)** Weight of the triceps surae muscle group (calf muscle) in affected and unaffected mice at three timepoints (30d, N = 17 WT, 7 Biltong; 9wk, and 6mo). **C)** Muscle weight to body weight ratio from mice taken at three timepoints (30d, 9wk, and 6mo). **D)** Average wirehang times of affected and unaffected mice at four timepoints (22d, 30d, 9wk, and 6mo). **E)** Nerve conduction velocities of affected and unaffected mice at four timepoints (22d, 30d, 9wk, and 6mo). **F-H)** Proximal and distal CMAP amplitudes and proximal to distal CMAP ratios taken at four timepoints (22d, 30d, 9wk, and 6mo). **I)** Data from repetitive nerve stimulation (RNS) on mice taken at 4 timepoints (22d, 30d, 9wk, and 6mo). Results are plotted as a ratio of CMAP amplitudes from the 10^th^ stimulus to the 1^st^ stimulus in a 10 Hz train. Sample sizes (N) for each group are indicated directly on the graphs in blue text. P-values for A-C and E-H were determined using Welch’s one-way ANOVA and Dunnett’s T3 Multiple Comparisons Test in GraphPad Prism. P-values for D and I were determined using a Kruskal-Wallis test with Dunn’s Multiple Comparison Test in GraphPad Prism. All values shown are mean ± SD. *=p<0.05, **=p<0.01, ***=p<0.001, ****=p<0.0001 for all graphs.

At 22 days, *Biltong* and wildtype (WT) mice were virtually indistinguishable by eye, with similar body weights, wire hang times, and no overt motor phenotype. However, beginning around 30 d, *Biltong* mice exhibited a progressive decline in body weight, muscle weight, muscle weight to body weight ratio, and ability to hang from an inverted wire grid for 60 seconds (Figure 2A-D). No differences in NCV or CMAP amplitudes were observed with stimulation at proximal (hip) or distal (ankle) sites between WT and affected mice at any timepoint, and CMAP amplitudes elicited at the hip and ankle were roughly equal, indicating there is no failure in action potential propagation, axon loss, or failure in synaptic transmission in peripheral motor nerves (Figure 2E-H). A possible defect in neuromuscular junction (NMJ) function was observed during RNS, with decrement in CMAP amplitude of the tenth stimulus in a train compared to the first at 22 and 30 days, but this was not seen at later ages (Figure 2I). Immunofluorescent labeling of NMJs revealed decreased innervation in the soleus and plantaris of Biltong mice at all timepoints, not just at the early timepoints where the RNS decrement was present (Figure 3 & Supp. Fig 1). No differences were seen in the number axons in the motor branch of the femoral nerve or the number of motor neurons in spinal cord sections of Biltong mice compared to WT at 6 mo, indicating a lack of motor neuron degeneration as would be seen in ALS (Supp Fig 2).

**Figure 3.**
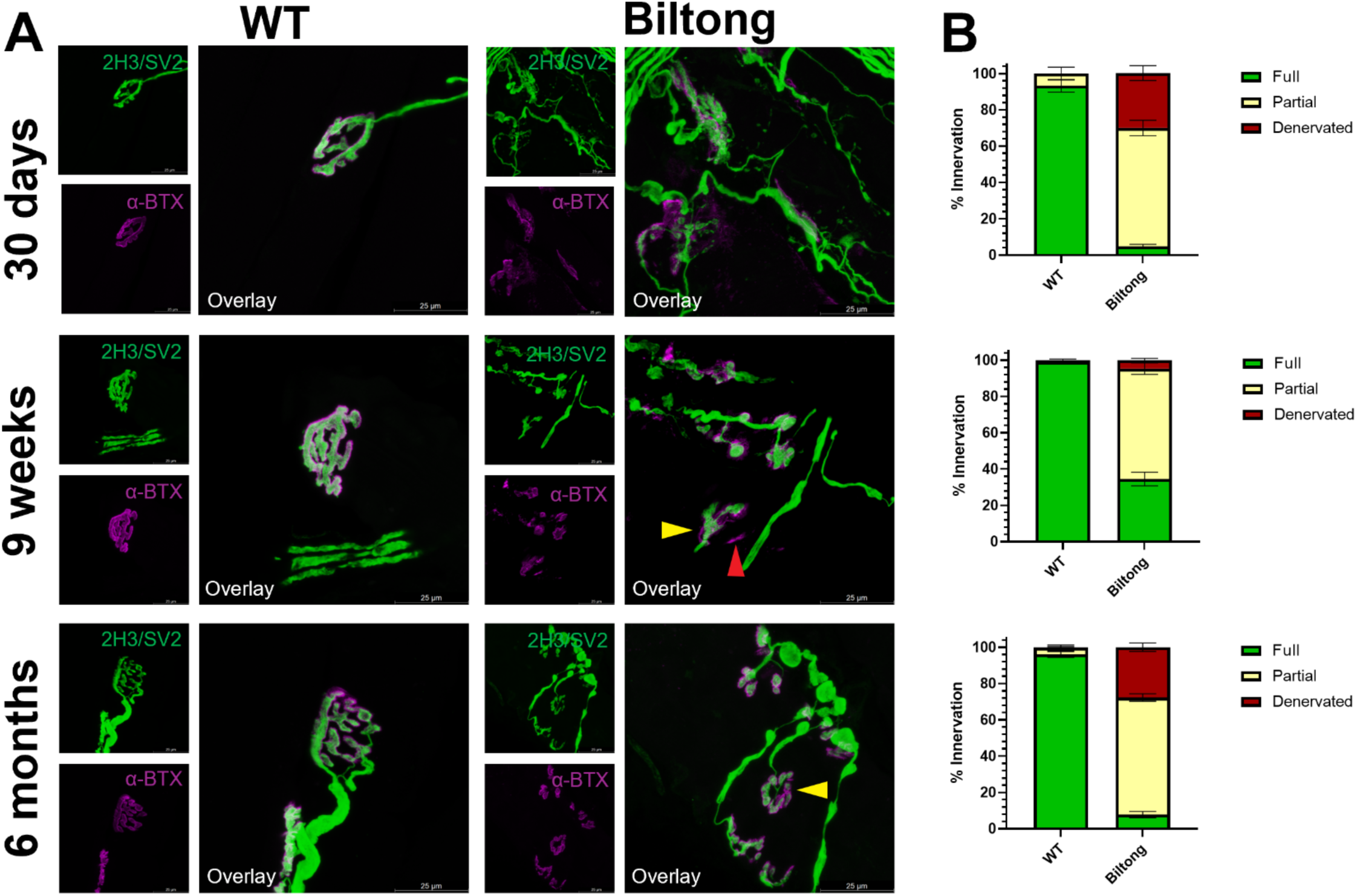
NMJ denervation in soleus of Biltong mice. **A)** Images of soleus NMJs from WT and Biltong mice at three timepoints (30d, 9wk, and 6mo). The axon terminal/presynapse is labeled in green with a cocktail of SV2 and NFM antibodies, while the acetylcholine receptors/postsynapse are labeled in magenta with AlexaFluor594-conjugated alpha-bungarotoxin. Yellow arrowheads in 9 wk and 6 mo Biltong images point to partially innervated NMJs and the red arrowhead in the 6 mo Biltong image points to a denervated endplate. Scale bars are 25 µm. **B)** Quantification of fully innervated (green), partially innervated (yellow), and denervated (red) NMJs from the same mice as in (A). N = 3 WT, 7 Biltong at 30 d; 10 WT, 11 Biltong at 9 wk; and 10 WT, 13 Biltong at 6 mo.

### Cerebellar Purkinje neuron degeneration

In addition to neuromuscular phenotypes such as muscle atrophy and decreased grip strength, Biltong mice also developed overt ataxia, which initially presented as an intention tremor when movements were initiated, suggesting a cerebellar phenotype. We therefore examined brains to look for pathological changes that could relate to the ataxia phenotype, including central demyelination and cerebellar dysfunction. Staining sagittal brain sections with luxol fast blue/crystal violet (LFB/CV) to visualize myelin did not reveal any major changes in brain myelination (not shown). Immunohistochemistry for calbindin to label cerebellar Purkinje neurons (PNs) showed normal cerebellar anatomy at 22 days (pre-onset) followed by rapid degeneration, with many PNs already lost by 30 days (Figure 4). PNs in lobule X were spared, even at 6 months of age when mice are approaching endpoint (Figure 4), indicating differential susceptibility/resilience to degeneration. A similar sparing of lobule X PNs has been reported in other ataxia mouse models^25,26^.

**Figure 4.**
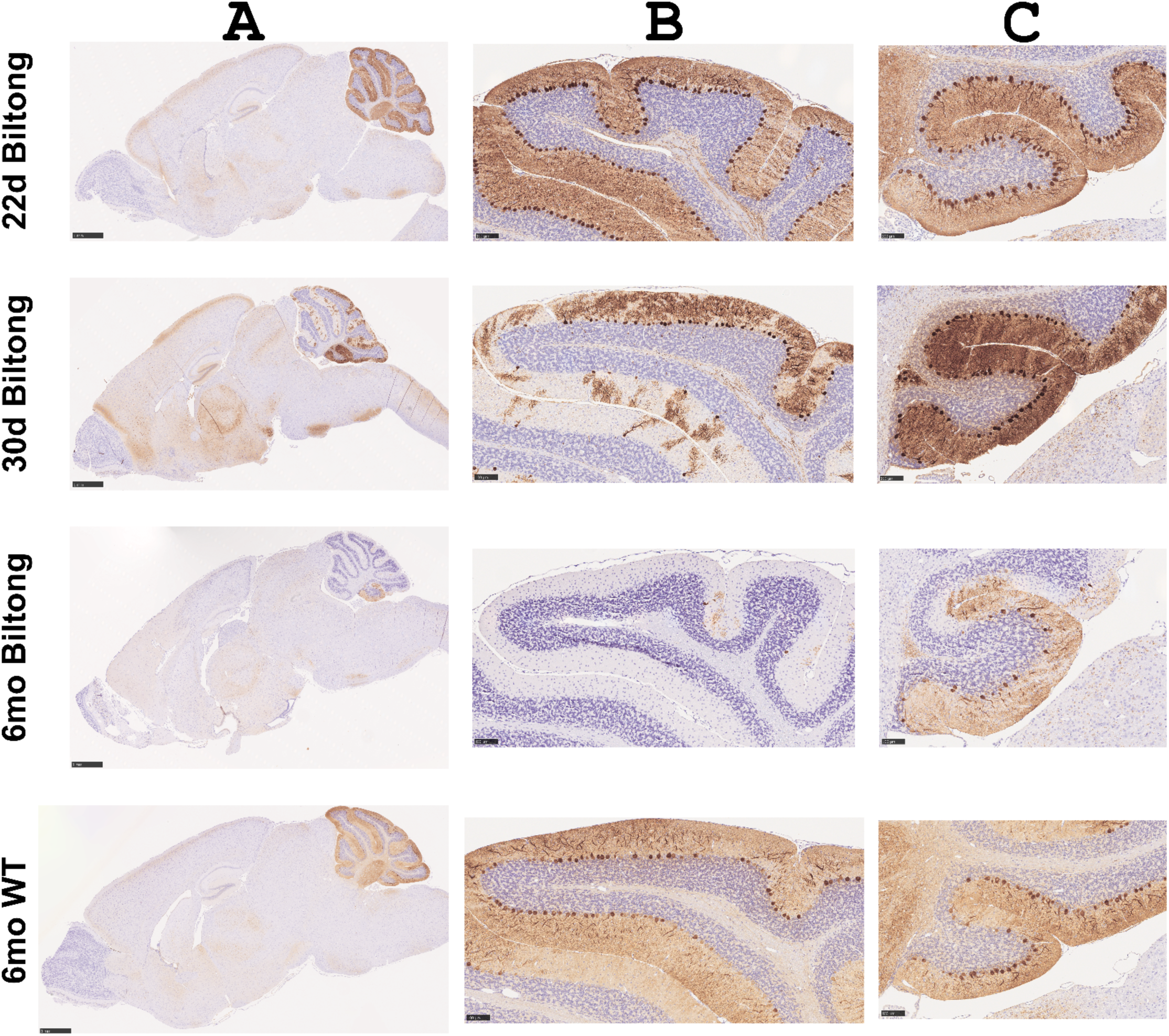
Calbindin staining reveals dramatic loss of Purkinje neurons in Biltong mice. **A)** Calbindin-stained sagittal brain sections taken from Biltong mice at three timepoints (22d, 30d, and 6mo) and from WT mice at 6mo. Scale bars are 1 mm. **B&C)** High magnification insets show lobules VI and X, respectively. Scale bars are 1 mm (whole brain) and 100 µm (lobules VI and X).

### Skeletal muscle pathology

Given the neuromuscular phenotypes described above and the emerging association of *TUBA4A* mutations with myopathy, muscles of the lower leg were examined by histopathology. Masson’s trichrome staining on cross sections of the lower hindlimb revealed fibrosis, central nuclei, inclusions, atrophic fibers, and ringbinden-like fibers (Figure 5). By 6 months, central nuclei, inclusions, and aggregates were observed in all muscles in the lower hindlimb, but other pathological changes were more spatially restricted. For example, fibrosis and atrophic/dystrophic fibers were most common in the soleus, while ringbinden-like fibers were found almost exclusively in the tibialis posterior. Similar to Purkinje cell degeneration, these defects developed around 30 days and progressed until 6 months.

**Figure 5.**
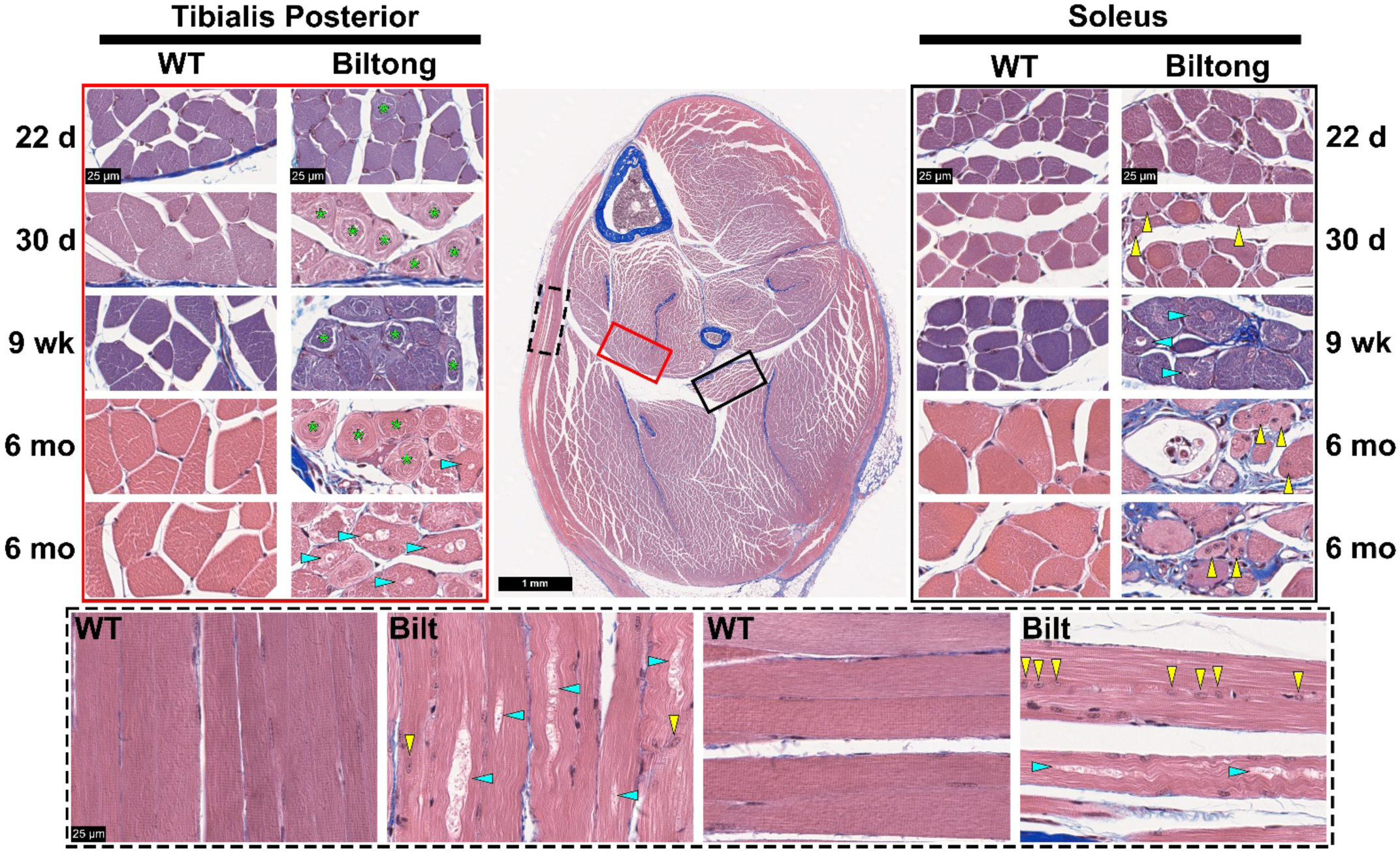
Skeletal muscle histopathology in Biltong mouse hindlimbs. Masson’s trichrome-stained lower hindlimb cross sections from WT and Biltong mice taken at four timepoints (22d, 30d, 9wk, and 6mo). High magnification (80x) images show the tibialis posterior (left, red rectangle), soleus (right, black rectangle), and longitudinal fibers from the hamstring (bottom, dashed black line, 6 month timepoint). Green asterisks denote ringbinden-like fibers, yellow arrowheads point to muscle fibers with centrally located nuclei, and cyan arrowheads point to fibers containing vacuoles. Scale bars = 1 mm (whole hindlimb section) or 25 µm (80x images).

Transmission electron micrographs (TEM) from cross-sections of the soleus show disrupted skeletal muscle architecture including disorganized myofibrils, Z-band streaming, multilamellar bodies, osmiophilic inclusions, central nuclei, and aberrant localization of mitochondria (Figure 6). Tibialis posterior was also imaged on TEM to view the ringbinden-like fibers seen in Trichrome staining. While numerous fibers with markedly disorganized myofibrillar structures were seen, none had the classic ringbinden cytoarchitecture seen in human myotonic dystrophies and myofibrillar myopathies^27–30^, in which myofibrils are arranged perpendicular to the cell membrane (Supplemental Figure 3).

**Figure 6.**
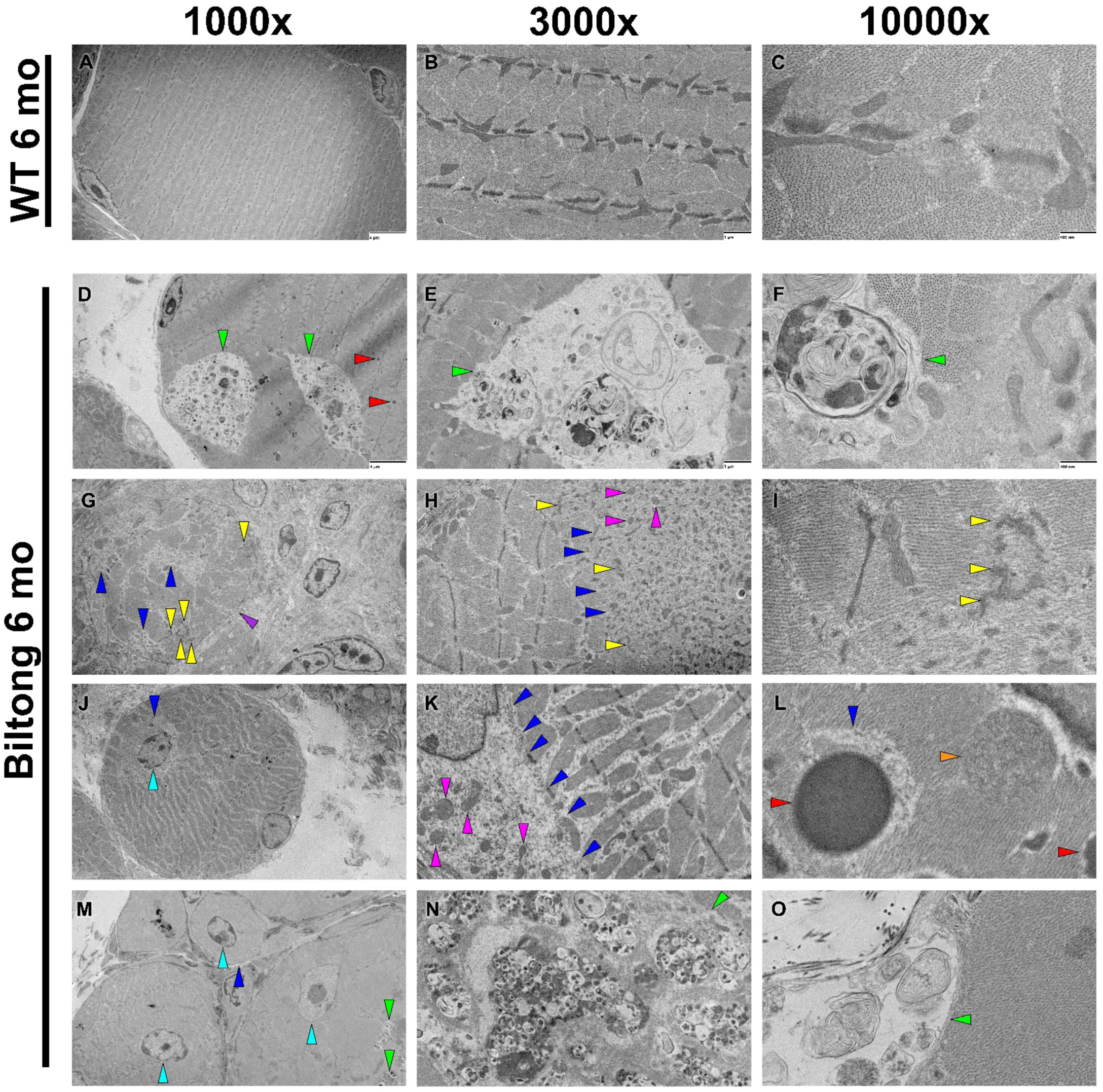
Biltong skeletal muscles have ultrastructural abnormalities. **A-C)** Representative transmission electron micrographs of cross-sectioned soleus muscle from WT mice at 6 months showing myofibers with normal ultrastructure. **D-O)** TEM images of cross-sectioned soleus from Biltong mice at 6 months show an array of ultrastructural abnormalities including regions containing osmiophilic inclusions (electron-dense spots) and multilamellar myeloid bodies (green arrowheads in D, E, F, M, N, and O), regions with disrupted myofibril organization (blue arrowheads in G, H, J, K, L, and M), Z-band streaming (yellow arrowheads in G, H, and I), centrally positioned myonuclei (cyan arrowheads in J and M), aberrant mitochondrial localization (magenta arrowheads in H and K), electron-dense cytoplasmic bodies (red arrowheads in D and L, orange arrowhead in L indicates a possible cytoplasmic body forming), atrophied myofibers (J and M), a degenerating muscle fiber (purple arrowhead in G), and fibrosis (right sides of G and J). Image magnification is denoted at the top of the figure. Scale bars are 4 µm (1000x), 1 µm (3000x), and 400 nm (10,000x) and are included in all WT images and the top row of Biltong images.

### *Tuba4a^Q176P^* recapitulates *Biltong* phenotypes, *Stk36^Y1003N^* does not

The analyses presented in figures 2-5 were performed in the Biltong mice identified from the mutagenesis screen and carrying mutations in both *Tuba4a* and *Stk36*. These variants were independently CRISPR-engineered into C57BL/6J (B6) mice which were then assessed for phenotypic changes such as jerky movements/ataxia, decreased body weight, skeletal muscle pathology, and degeneration of cerebellar Purkinje neurons. *Stk36^Y1003N^* mice (C57BL/6J-Stk36<em1Rwb</Rwb) did not develop any of these phenotypes, even at 6 months of age when the Biltong mice are near end-stage (Supplemental Figure 4). However, the *Tuba4a^Q176P^* mice (CByJ;B6J -Tuba4a<em1Rwb>/Rwb) reproduced all phenotypes at 9 weeks (Figure 7). Conventional breeding of *Tuba4a^Q176P^* founders to wild-type B6 mice produced only a single litter and these offspring did not breed. Therefore, an IVF to BALB/cByJ was performed to rescue the line and the mice were maintained on a segregating B6/BALB background. Sperm agglutination was noticed when taken for IVF, which could relate to the difficulties breeding. A similar phenomenon was observed in the original *Biltong* strain – as they were backcrossed to B6 or BALB, they produced fewer litters and eventually stopped breeding.

**Figure 7.**
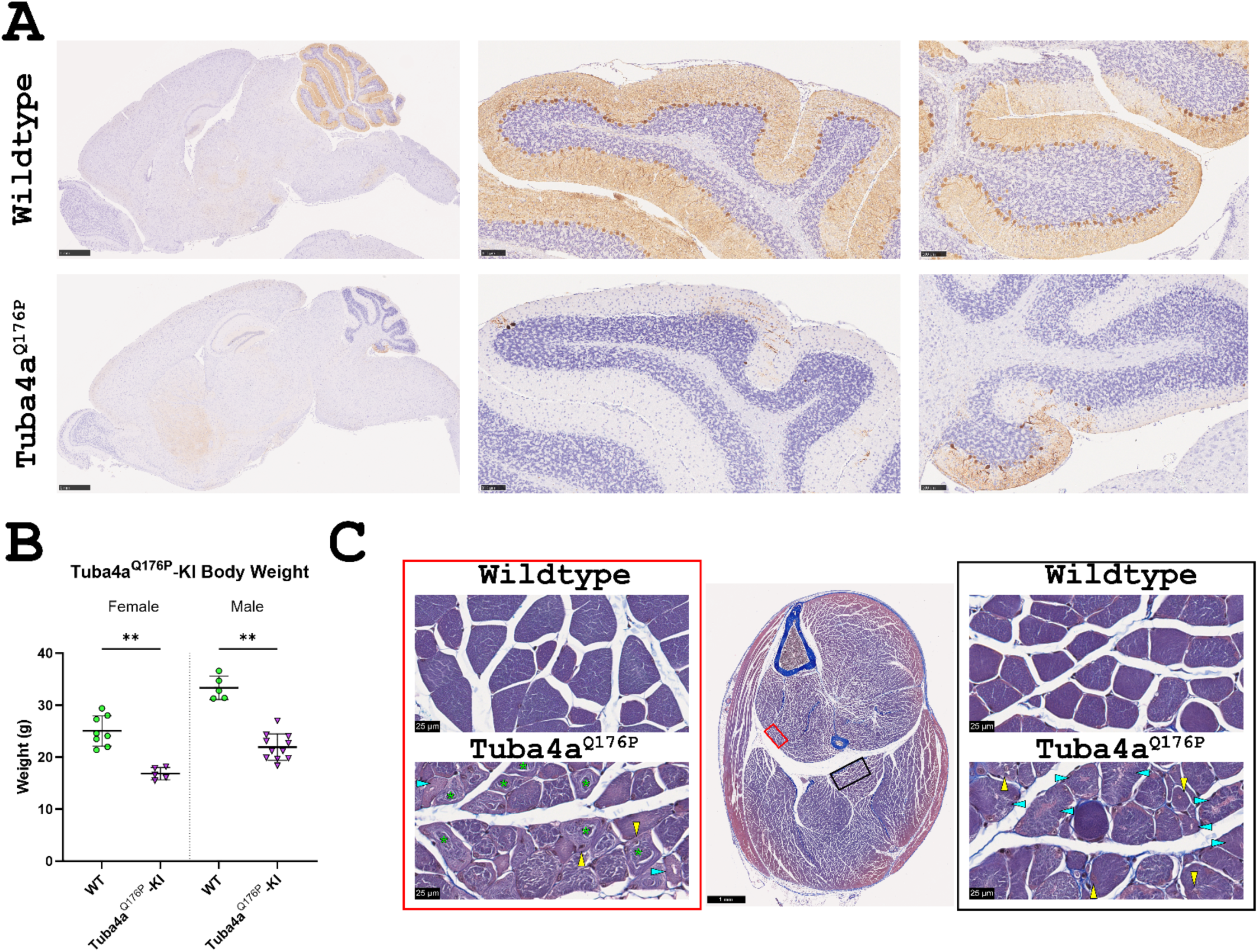
Knockin of *Tuba4a^Q176P^* reproduces Biltong phenotypes by 9 weeks. **A)** Sagittal brain sections from 9 wk WT and *Tuba4a^Q176P^* knockin mice stained with Calbindin to label cerebellar Purkinje neurons. Scale bars are 1 mm (whole brain) and 100 µm (lobules VI and X). **B)** Body weights from female and male mice at 9 wk. **p < 0.01 **C)** Masson’s trichrome staining of lower hindlimb cross-sections from WT and *Tuba4a^Q176P^* knockin mice taken at 9 wk. High magnification (80x) images show the tibialis posterior (left, red rectangle), and soleus (right, black rectangle). Green asterisks denote ringbinden-like fibers, yellow arrowheads point to muscle fibers with centrally located nuclei, and cyan arrowheads point to fibers containing vacuoles. Scale bars = 1 mm (whole hindlimb section) or 25 µm (80x images).

## DISCUSSION

We identified mice with an overt neuromuscular/ataxia phenotype in an ENU mutagenesis screen and subsequent breeding was consistent with a fully penetrant dominant mode of inheritance. Phenotyping confirmed defects in neuromuscular innervation, extensive muscle histopathology, and cerebellar Purkinje neuron degeneration, all beginning around four weeks of age. Genetic mapping and genome sequencing revealed candidate mutations in *Tuba4a* and *Stk36*. The causality of the *Tuba4a*^Q176P^ mutation was confirmed by engineering this variant into wild type mice, which reproduced the phenotype of the mice from the ENU screen, whereas mice with the *Stk36*^Y1003N^ mutation were unaffected. In addition, Western blot of lysates from skeletal muscle and cerebellum of Biltong mice (from the mutagenesis) showed reduced TUBA4A protein levels.

Dominant mutations in alpha tubulin 4A (*TUBA4A*) are associated with spastic ataxia type 11 (SPAX11), congenital myopathy 26 (CMYO26), and frontotemporal dementia and amyotrophic lateral sclerosis 9 (FTDALS9) in humans^8–12,16–20^. The *Tuba4a*^Q176P^ mice have muscle pathology that is remarkably similar to patients with congenital myopathy caused by a *de novo* mutation (L227F) in *TUBA4A*, which causes “rimmed vacuoles, cytoplasmic TUBA4A aggregates, and myofibrillar disorganization” with infantile onset^19^. A subsequent report identified more variable clinical myopathy presentations and modes of inheritance associated with *TUBA4A* mutations^18^. However, the *Tuba4a*^Q176P^ mouse appears to be a valid model for at least some *TUBA4A*-associated myopathy patients.

The *Tuba4a*^Q176P^ mice have profound ataxia with Purkinje neuron degeneration beginning between 22 and 30 days. Ataxia and cerebellar atrophy were also observed in two myopathy patients from the previously mentioned study^16^. SPAX11 has a highly variable age of onset and may include other features such as dysmetria and mild sensorimotor neuropathy. Some cerebellar abnormalities including possibly enlarged interfolial spaces in the crus were observed by MRI in one patient^17^. The overall anatomy of the cerebellum is preserved in the *Tuba4a*^Q176P^ mice, but nearly all Purkinje cells are eventually lost except in the most caudal cerebellar lobule. Whether the mice have spasticity remains to be examined. There are markers of upper motor neurons in the cortex and axons in the corticospinal tract in the spinal cord, but careful quantification and matching is required and has not been undertaken at this time.

We did not find phenotypes consistent with motor neuron disease in the *Tuba4a*^Q176P^ mice. Lower motor neurons in the spinal cord were preserved and there was no reduction in peripheral motor axon number or changes in neurophysiology such as reduced CMAP amplitude that would be consistent with motor neuron loss. In patients, the clinical FTDALS9 presentation is quite variable, with some patients showing Frontotemporal Dementia (typically behavioral variant FTD), whereas others show ALS, and some present with both^8–12,20^. The onset is typically in the sixth or seventh decade of life, and thus may be later than our latest time point (six months) in the mouse studies. It is possible that FTDALS9-associated mutations impact upper and lower motor neurons more severely due to distinct mechanisms from those that cause SPAX11 or CMYO26. It is also possible that with more time, the *Tuba4a*^Q176P^ mice would develop motor neuron disease, but testing this may require a conditional approach to prevent welfare concerns after 6 months of age. In some FTDALS9 patients, TDP43 pathology was observed, but this did not colocalize with tubulin aggregates, raising the question of whether it is a secondary effect of ongoing neurodegeneration. We have not examined TDP43 or tubulin aggregates in the mice.

The motor deficits seen in SPAX11 patients are clinically distinct from those seen in FTDALS9, as there is no lower motor neuron involvement^16^. However, cognitive impairment has been reported in a subset of SPAX11 cases (5 of 24 patients across two studies), indicating a potential link between these disorders^16,17^. To further support this, the *TUBA4A^R105C^*variant has been identified both in a case of SPAX11 and in a large, multi-generational Dutch family containing seven individuals with dementia (3 with FTD and 4 with unspecified dementia, UD)^9,16^. The G11H mutation was identified in a patient who presented with ataxia and myofibrillar myopathy and received a diagnosis of multisystem proteinopathy^18^. There also appears to be a connection to Parkinson’s disease (PD): one member of the previously mentioned family with UD also had parkinsonism and another was diagnosed PD^9^; another FTDALS9-associated mutation (R64Gfs90*) was found in a patient with semantic dementia whose father and paternal grandmother were diagnosed with PD^10^; and one study reported a case of “pure nigropathy” (without Lewy pathology) associated with a nonsense mutation in *TUBA4A* (R79X)^31^. These observations suggest that *TUBA4A*-related disorders exist on a clinical and genetic spectrum encompassing spastic ataxia, FTD, ALS, and possibly PD.

*Biltong* mice were poor breeders (few litters and shortened lifespan), and this was more pronounced on an inbred B6 or BALB genetic background. However, the mutation does not impact offspring viability since affected mice were born at the expected Mendelian ratio (43 affected mice of 91, Χ^2^ = 0.275, df = 1, p = 0.6002). *TUBA4A* mutations are also associated with female infertility and early zygotic arrest in humans^21–23^. We have not explored the basis for the reduced fertility, but anecdotally, males were also poor breeders and IVF using male sperm donors was required to propagate both the ENU and CRISPR allele of *Tuba4a*. Given the severe and early ataxia, the issue may be behavioral and not reproductive. Female fertility could be tested by ovary transplantation from a mutant female to a compatible host, but eliminating effects of surgical technique makes this a challenging experiment to do quantitatively.

The fully penetrant dominant phenotype of the *Tuba4a*^Q176P^ mutation is consistent with a dominant negative/toxic gain-of-function effect of the mutation. Mice with a complete knockout allele of *Tuba4a* have been reported and do not have a notable phenotype unless combined with knockout of beta tubulin 1 (*Tubb1*), which causes impaired hemostasis and spherocytosis, a disorder in which platelets exhibit a spherical shape rather than their typical discoid shape^32^. In the process of CRISPR engineering the confirmatory Q176P allele, we also generated a 29 base pair deletion in *Tuba4a*, resulting in a frame shift and likely loss-of-function allele (CByJ;B6J-Tuba4a<em2Rwb>/Rwb). These mice do not have an overt phenotype as heterozygotes, arguing against haploinsufficiency. Since the dominant Q176P mutation has a more severe phenotype than the complete knockout, presumably it is interfering with tubulins more generally. Given the dimerizing and polymerizing nature of tubulins, such an effect is plausible. Homozygous Q176P mice are not viable postnatally, indicating an even more severe impact of the mutation as gene dosage increases. We have not explored the embryonic phenotype of these mice to know if this constitutes a zygotic/embryonic arrest reported in humans. The juvenile onset of the phenotypes in the mice may be explained by a tissue- or cell type-specific developmental switch in alpha tubulin gene expression to *Tuba4a* around the time of onset. This idea is supported by data showing low *Tuba4a* mRNA levels in the developing mouse brain that dramatically increase by 8 weeks^33^.

In summary, the *Tuba4a* allele identified in this ENU mutagenesis screen (based on its overt ataxia phenotype) appears to be a valid model of both human ataxias and myopathies with both appropriate genetics and relevant phenotypes^34^. Whether the mice also model female infertility and upper motor neuron pathology consistent with FTDALS9 will require further examination. These mice will be useful for understanding pathophysiology caused by tubulin dysfunction in both neurons and skeletal muscle. They also provide a good animal model for optimizing gene therapy approaches such as allele-specific knockdown or CRISPR-mediated editing to correct or silence the mutant allele that may apply to *TUBA4A* mutations and other tubulinopathies.

## METHODS

### Generation and maintenance of the ENU-induced mutant line

Mutagenesis using N-ethyl-N-nitrosourea (ENU) was performed as previously described^35–37^. Briefly, male C57BL6/J mice were injected with 80 mg/kg ENU three times at weekly intervals and mated with C57BL6/J female mice to produce G1 offspring, which were monitored for development of potential neurological phenotypes. A G1 male developed ataxia around 4 weeks of age and IVF was performed with BALB/cByJ (BALB) oocyte donors to propagate the line. Affected F1 offspring were crossed again to WT BALB to generate N2 mice for mapping, followed by further backcrosses to both B6 and BALB to make congenic strains. However, the mice stopped breeding after 5-6 backcrosses to either strain, so they were maintained on a mixed B6 x BALB background (official strain name: CByJ;B6J-Tuba4a<Bilt>/Rwb). All mice were maintained in the same vivarium on a 14:10 light/dark cycle and were provided food and water ad libitum. Care and procedures were reviewed and approved by the Animal Care and Use Committee of The Jackson Laboratory.

### Genetic mapping and whole exome and genome sequencing

All genomic coordinates used in this manuscript refer to NCBI Mouse Genome build 38 (GRCm38/mm10). Genomic DNA was isolated from tail snips of 22 affected and 22 unaffected N2 mice for genetic mapping. A genotyping panel containing 142 genome-wide single nucleotide polymorphisms (SNPs) that differ between B6 and BALB strains was run on these DNA samples. From this panel, 1 SNP on chromosome 1 (rs3677697, located at 80,181,487 bp) segregated almost perfectly with the affected animals (one unaffected animal was also positive for this SNP. The flanking SNPs (rs3673812, at 62,173,915 bp and rs13476023 at 103,636,838 bp) did not segregate with the affected mice (6 and 4 mismatches, respectively). Additional mapping was performed using 8 B6/BALB SNPs within the ∼41 Mbp interval. From this panel, two SNPs (rs3672814 at 78,030,398 bp and rs4222476 at 79,289,860 bp) segregated perfectly with affected mice, while the flanking SNPs (rs3683684 at 69,020,216 bp and rs3664528 at 94,081,544 bp) had 1 and 3 mismatches, respectively (Figure 1A). This defined a ∼25Mbp region of interest from 69,020,216 – 94,081,544 bp on chromosome 1. A list of genes in this interval can be found in Supplemental Table 1.

For whole exome sequencing (WES), genomic DNA was isolated from spleens of two affected mice and one unaffected mouse using the Wizard DNA Purification Kit (Promega) according to the manufacturer’s protocols. DNA concentration and quality were assessed using the Nanodrop 2000 spectrophotometer (Thermo Scientific), the Qubit 3.0 dsDNA BR Assay (Thermo Scientific), and the Genomic DNA ScreenTape Analysis Assay (Agilent Technologies). Mouse exome libraries were constructed using the KAPA HyperPrep Kit (Roche Sequencing and Life Science) and SureSelectXT Mouse All Exon V2 Target Enrichment System (Agilent Technologies), according to the manufactures’ protocols. Briefly, the protocol entails shearing the DNA using the E220 Focused-ultrasonicator (Covaris) targeting 200 bp, ligating Illumina-specific adapters, and PCR amplification. The amplified DNA libraries are then hybridized to the Mouse All Exon probes (Agilent Technologies) and amplified using indexed primers. The quality and concentration of the libraries were assessed using the High Sensitivity D5000 ScreenTape (Agilent Technologies) and Qubit dsDNA HS Assay (ThermoFisher), respectively, according to the manufacturers’ instructions. Libraries were sequenced 75 bp paired-end on an Illumina NextSeq 500 using the High Output Reagent Kit v2.5. This method revealed a coding SNP in *Stk36* that segregated with the ataxia phenotype.

When no obvious phenotype was observed after CRISPR engineering the *Stk36* variant into wild-type mice, we performed whole genome sequencing (WGS) on genomic DNA extracted from spleens of three affected mice and one unaffected mouse. DNA was isolated from tissue using the NucleoMag Tissue Kit (Machery-Nagel) according to the manufacturer’s protocol. DNA concentration and quality were assessed as described above. Whole genome libraries were constructed using the KAPA HyperPrep Kit according to the manufacturer’s protocols, targeting an insert size of 400 base pairs. Briefly, the protocol entails shearing the DNA using the E220 Focused-ultrasonicator, size selection targeting 400 bp, ligating Illumina-specific barcoded adapters, and PCR amplification. The quality and concentration of the libraries were assessed using the methods described above. Libraries were sequenced 150 bp paired-end on an Illumina NovaSeq X Plus using the 10B Reagent Kit. This analysis identified the mutation in *Tuba4a*, which was present in the WES, but in a region of low read depth and therefore not initially identified as a candidate.

Fastq files from both experiments were processed using a pipeline developed at The Jackson Laboratory, which filters out known SNPs between commonly used mouse strains. DNA sequences were then viewed in Integrated Genome Viewer (IGV) to identify potential causative mutations.

### Genotyping

Ear notches from mice were digested in Proteinase K overnight at 55°C, then Proteinase K was heat inactivated by boiling for 10 min. PCR was performed using the primers below, and PCR products were sent to JAX’s Genome Technologies service for Sanger sequencing. Sequences were evaluated using Technelysium Chromas chromatogram viewer.

The following primers were used to amplify a 446bp segment of DNA containing the segregating *Stk36* mutation:

> Fwd – 5’ ACACAGTGGCCACCTCTTCT 3’
>
> Rev – 5’ AAGCAGCTGGGTCATACTGG 3’

The following primers were used to amplify a 913bp segement of DNA containing the causative *Tuba4a* mutation:

> Fwd – 5’ AGCCTACTGTAATCGGTGAGCA 3’
>
> Rev – 5’ TATCTAGGTTACGGCGGCAGA 3’

### Western Blot

Mice were euthanized at 6 months by CO_2_ asphyxiation and the cerebellum and soleus muscles were harvested and snap frozen for downstream Western blot analysis. Tissues were dounce homogenized in RIPA buffer with 1X Halt Protease and Phosphatase Inhibitor Cocktail (ThermoScientific) and protein concentration was measured using the Pierce BCA Protein Assay Kit (ThermoScientific) according to manufacturer instructions. 12 µg total protein was run in each well of a polyacrylamide gel, then transferred to polyvinylidene difluoride (PVDF) membrane. The membrane was blocked with 5% non-fat dry milk dissolved in 1X phosphate buffered saline with 0.1% Tween-20 (PBS-T) for 1 h, then incubated in primary antibodies overnight at 4°C. HRP-conjugated secondary antibodies were diluted in the same blocking solution and incubated with the membrane for 1 h at room temperature, then bands were visualized using Super Signal West Pico Plus Chemiluminescent Substrate (ThermoScientific, 34580) and X-ray film (Thomas Scientific, 1141J52). Films were scanned and band intensity was measured using ImageJ.

Primary antibodies used were rabbit anti-TUBA4A (1:1000, abcam, ab177479) and mouse anti-alpha tubulin (1:3000, Cell Signaling Technologies, 3873T). Secondary antibodies used were HRP-conjugated donkey anti-rabbit (1:25,000, Jackson Immunoresearch, 711-035-152) and HRP-conjugated horse anti-mouse (1:5000, Cell Signaling Technologies, 7076P2).

### Electromyography

To measure nerve conduction velocity (NCV) and response to repetitive nerve stimulation (RNS), electromyography (EMG) was performed as previously described^38^. For NCV, mice were anesthetized with 3% isoflurane on a heating pad to maintain body temperature. Needle recording electrodes were placed as such: positive (+) in the plantar muscle of the hindpaw on the same side as the stimulus, negative (-) in the fifth toe of that same foot, and ground in the plantar muscle of the opposite hindpaw. Teflon-coated stimulating electrodes were inserted subcutaneously adjacent to the sciatic nerve, first at the ankle, then at the sciatic notch. The nerve was then stimulated with 1-3 mV and compound muscle action potential (CMAP) amplitude was recorded and used to calculate NCV.

For repetitive nerve stimulation (RNS), mice were anesthetized as described above and recording electrodes were placed in the left hindpaw (ground), the tibalis anterior (negative), and the right hindpaw (positive). Stimulating electrodes were placed at the sciatic notch in the hip. Stimulating voltage was increased until maximal CMAP amplitude was achieved. Consecutive 10-pulse trains were then administered at the following frequencies: 3 Hz, 6 Hz, 10 Hz, 20 Hz, 50 Hz, 60 Hz, and 75 Hz. To mitigate the effects of repeated trains, mice were allowed to rest for five seconds between trains up to 20 Hz and for ten seconds between trains of higher frequencies. Stimulation voltage during the trains was applied as 125% of the voltage needed to elicit maximal CMAP amplitude during baseline. Decrement was calculated by dividing the CMAP amplitude elicited by the 10^th^ pulse of a train to the CMAP amplitude elicited by the first pulse. Data shown in Figure 2 are from the 10 Hz train, but a similar pattern was seen at all frequencies.

### Inverted wire hang

To test motor function, the inverted wire hang test was used as previously described^38^. Briefly, mice were suspended from an inverted wire grid for a maximum trial time of 60 s. Three trials were performed on each mouse at each timepoint and the average hang time of the three trials is reported. Mice were not trained prior to the initial test or in between timepoints.

### Immunohistochemistry

For calbindin immunohistochemistry (IHC) to label cerebellar Purkinje neurons in the brain, mice were terminally anesthetized with a lethal dose of tribromoethanol, then cardiac perfusion was performed with 1X phosphate buffered saline (PBS) followed by 10% neutral buffered formalin (NBF). Brains were removed, cut in half at the midline and immersion-fixed in NBF overnight at 4°C. Brains were paraffin-embedded, sectioned sagittally at 5 μm, and IHC for calbindin (ab108404, abcam, 1:1000) was performed using the Bond RX Automated Research Stainer (Leica Biosystems) with the Bond Polymer Refine Detection Kit (DS9800, Leica Biosystems).

For ChAT IHC to label motor neurons in the spinal cord, mice were euthanized with CO_2_ and decapitated. Spinal cords were removed using hydraulic extrusion and immersion-fixed in 10% NBF overnight at 4°C. Cords were then paraffin-embedded, cross-sectioned at 5 μm, and IHC for ChAT (ab178850, abcam, 1:2000) was performed with the Bond RX platform as described above.

### Skeletal muscle histology

After euthanasia by CO_2_ asphyxiation, mice were skinned and whole hindlimbs were removed and immersion-fixed in Bouins fixative for 1 week to decalcify bones. The tissue was trimmed and a 2-3 mm cross-section of the mid-calf was paraffin-embedded, sectioned at 4 μm, and processed for Trichrome staining (Masson’s method), which was performed manually.

### NMJ staining

Triceps surae were isolated and weighed for muscle weight to body weight ratios, then the soleus and plantaris were removed and fixed for 1-2 h in fresh, ice-cold 2% paraformaldehyde (PFA) diluted from ampules of 16% PFA (Electron Microscopy Services) in 1X PBS. Muscles were then shredded using fine forceps and blocked with 2% bovine serum albumin (BSA) block (2% BSA + 0.2% Triton X-100 in 1X PBS) while being compressed between two microscope slides that were held together with binder clips. The axon terminal (presynapse) was labeled with a cocktail of mouse anti-NEFM (1:500, 2H3-c, Developmental Studies Hybridoma Bank (DSHB)) and mouse anti-SV2 (1:250, SV2-c, DSHB), followed by incubation with an AlexaFluor488-conjugated anti-mouse secondary antibody and AlexaFluor594-conjugated α-bungarotoxin (to label postsynaptic acetylcholine receptors). NMJs were imaged using a Leica Stellaris 8 WLL confocal microscope. Fully innervated NMJs were those in which the presynaptic terminal completely covered the postsynaptic receptor field. In partially innervated NMJs, there were regions of the postsynaptic receptor field that were not in apposition to the presynaptic terminal. In denervated NMJs, sites of AChR labeling no longer had associated presynaptic structures.

### Transmission electron microscopy (TEM)

Motor and sensory branches of the femoral nerve, as well as soleus and tibialis posterior (TP) muscles were dissected and immersion-fixed in EM fixative (4% PFA, 2.5% glutaraldehyde, in cacodylate buffer) overnight, then plastic-embedded. For nerves, semithin (0.5 μm) sections were stained with toluidine blue to visualize myelin. These sections were imaged at 40X magnification on a Nikon Eclipse E600 microscope with a QIClick Mono 12-bit camera (Q-Imaging). For muscles, ultrathin cross-sections were mounted on TEM grids and imaged on a JEOL JEM-1230 TEM with an AMT NanoSprint15 Mk-II camera.

### CRISPR-engineered Stk36 and Tuba4a mice

*Stk36^Y1003N^*and *Tuba4a^Q176P^*mutant mice were generated on a C57BL/6J background using CRISPR/Cas9 and maintained on a mixed B6/BALB genetic background for breeding purposes (official strain names: C57BL/6J-Stk36<em1Rwb>/Rwb and CByJ;B6J -Tuba4a<em1Rwb>/Rwb, respectively.

Two overlapping guide RNAs were used to target Cas9 to the *Stk36* gene:

**Guide 1** – ACTGTGCGGGTCCGTGCATA

**Guide 2** – CTGTGCGGGTCCGTGCATAT

The sequence of the Donor DNA used to generate the mutant *Stk36* allele (**silent** and **of interest** mutations): TAGCAGCAGAAGGTTGAGCAGTCCTGACTGACTCTGTAGTGCCCCACGTAGGGCCATGCTGTGCTGTAAT AAATGTCCCAGGAG**A**CCAT**TA**GC**T**CGGAC**G**CGCACAGTGTTCTCTGAGTGGCCTAGG

Two overlapping guide RNAs were used to target Cas9 to the *Tuba4a* gene locus:

**Guide 1** – CACTACAGCCGTGGACACTT

**Guide 2** – TACAGCCGTGGACACTTGGG

The sequence of the Donor DNA used to generate the *Tuba4a* allele (**causative mutation**): GTCTGAGTGTTCCAGGGTGGTATGGGTGGTCAGGATGGAGTTGTAGGGCTCCACTACAGCCGTGGACACT **G**GGGGGGCTGGGTAGATGGAGAATTCCAGCTTGGACTTTTTGCCATAGTCAACAGAAAGCCGCTCCATCA G

### Statistics

#### Routine statistics

Statistical analyses were performed in GraphPad Prism 10. Data was not tested for normality, as sample sizes were generally not large enough to be meaningful. No outliers were excluded and all relevant data points were included in the analyses. The specific statistical tests used are mentioned in the Results and corresponding Figure captions.

#### Chi-square analysis

Chi-square analysis was performed using the GraphPad ChiSquared Calculator online (https://www.graphpad.com/quickcalcs/chisquared1/) using data from all full litters of mice that were phenotyped or had tissue collected at the 22 and 30 day timepoints. Pre-onset mice were genotyped using primers for *Tuba4a* that amplify the region containing the causative mutation, followed by Sanger sequencing.

Post-onset mice were initially grouped based on presence of the phenotype, then confirmed with Tuba4a genotyping (which did not alter the groupings). 43 affected and 48 unaffected mice were observed from 91 total mice, giving a chi-squared (Χ^2^) value of 0.275 with a p-value of 0.6002.

## ACKNOWLEDGEMENTS

We gratefully acknowledge the contribution of the Scientific Services at The Jackson Laboratory, particularly Genome Technologies, Genetic Engineering Technologies, Reproductive Sciences, Computational Sciences, Histopathology, and Electron Microscopy services for expert assistance with the work described in this publication. We also thank Louise Dionne at The Jackson Laboratory for bringing the original mutagenized male to our attention.

## Abbreviations

ENU: N-ethyl-N-nitrosourea
SNP: single nucleotide polymorphism
CMYO: congenital myopathy
SPAX: spastic ataxia
FTD: frontotemporal dementia
ALS: amyotrophic lateral sclerosis
CRISPR: clustered regularly interspaced short palindromic repeats
Bp: base pairs
WES: whole exome sequencing
WGS: whole genome sequencing
EMG: electromyography
CMAP: compound muscle action potential
NCV: nerve conduction velocity
RNS: repetitive nerve stimulation
WT: wild-type
NMJ: neuromuscular junction
CV/LFB: cresyl violet luxol fast blue
PN: Purkinje neuron
TEM: transmission electron microscop
IVF: in vitro fertilization
MRI: magnetic resonance imaging
PD: Parkinson’s disease
PCR: polymerase chain reaction
DNA: deoxyribonucleic acid
PVDF: polyvinylidene difluoride
PBS-T: phosphate buffered saline with Tween 20
HRP: horseradish peroxidase
NBF: neutral buffered formaline
PFA: paraformaldehyde
BSA: bovine serum albumin

---

**Supplementary Figure 1.**
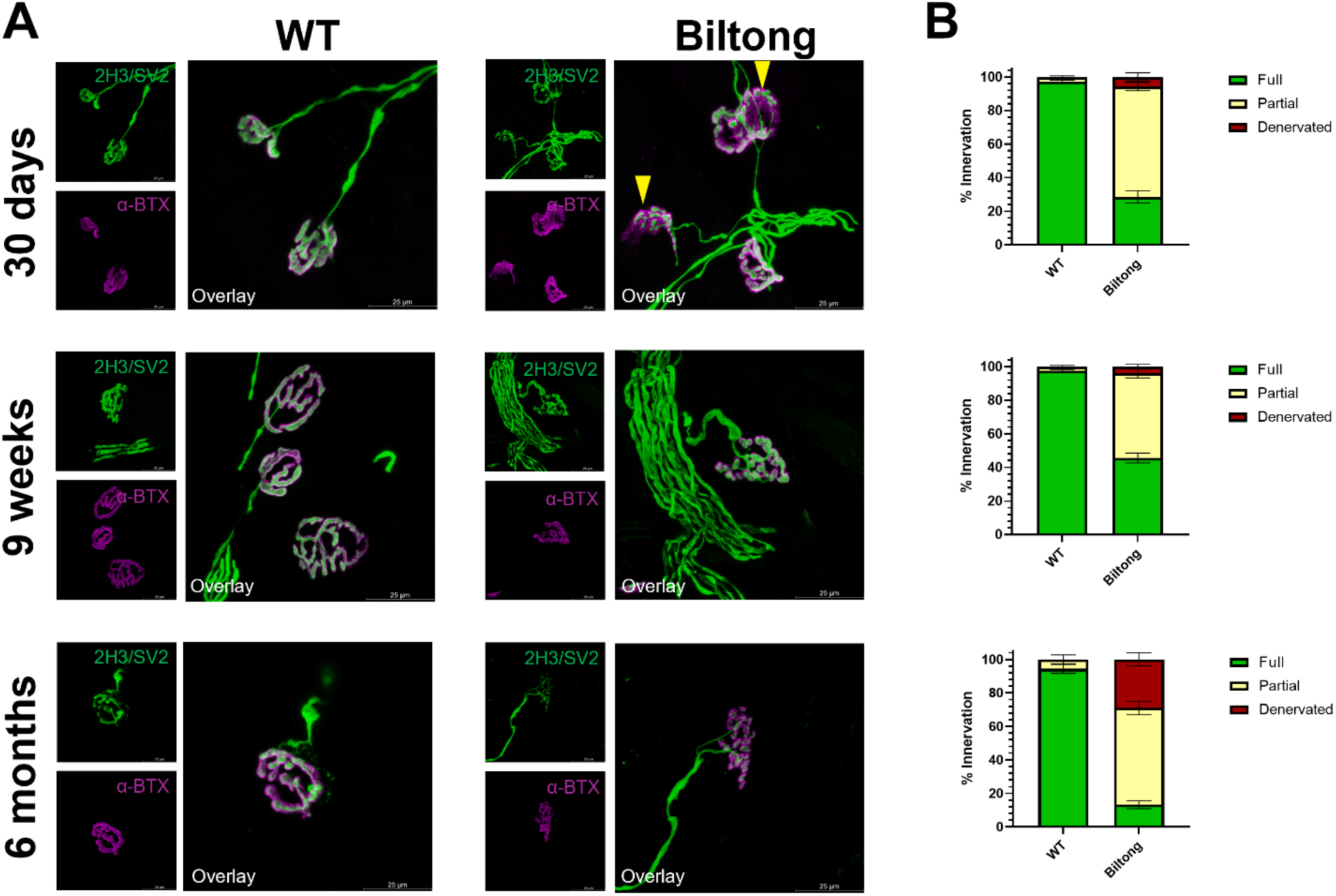
NMJ denervation in plantaris of Biltong mice. **A)** Images of plantaris NMJs from WT and Biltong mice at three timepoints (30d, 9wk, and 6mo). The axon terminal/presynapse is labeled in green with a cocktail of SV2 and NFM antibodies, while the acetylcholine receptors/postsynapse are labeled in magenta with AlexaFluor594-conjugated α-bungarotoxin. Yellow arrowheads in 30 d Biltong image indicate partially innervated NMJs. Scale bars are 25 μm. **B)** Quantification of fully innervated (green), partially innervated (yellow), and denervated (red) NMJs from the same mice as in (A). N = 3 WT, 7 Biltong at 30 d; 10 WT, 11 Biltong at 9 wk; and 10 WT, 13 Biltong at 6 mo.

**Supplementary Figure 2.**
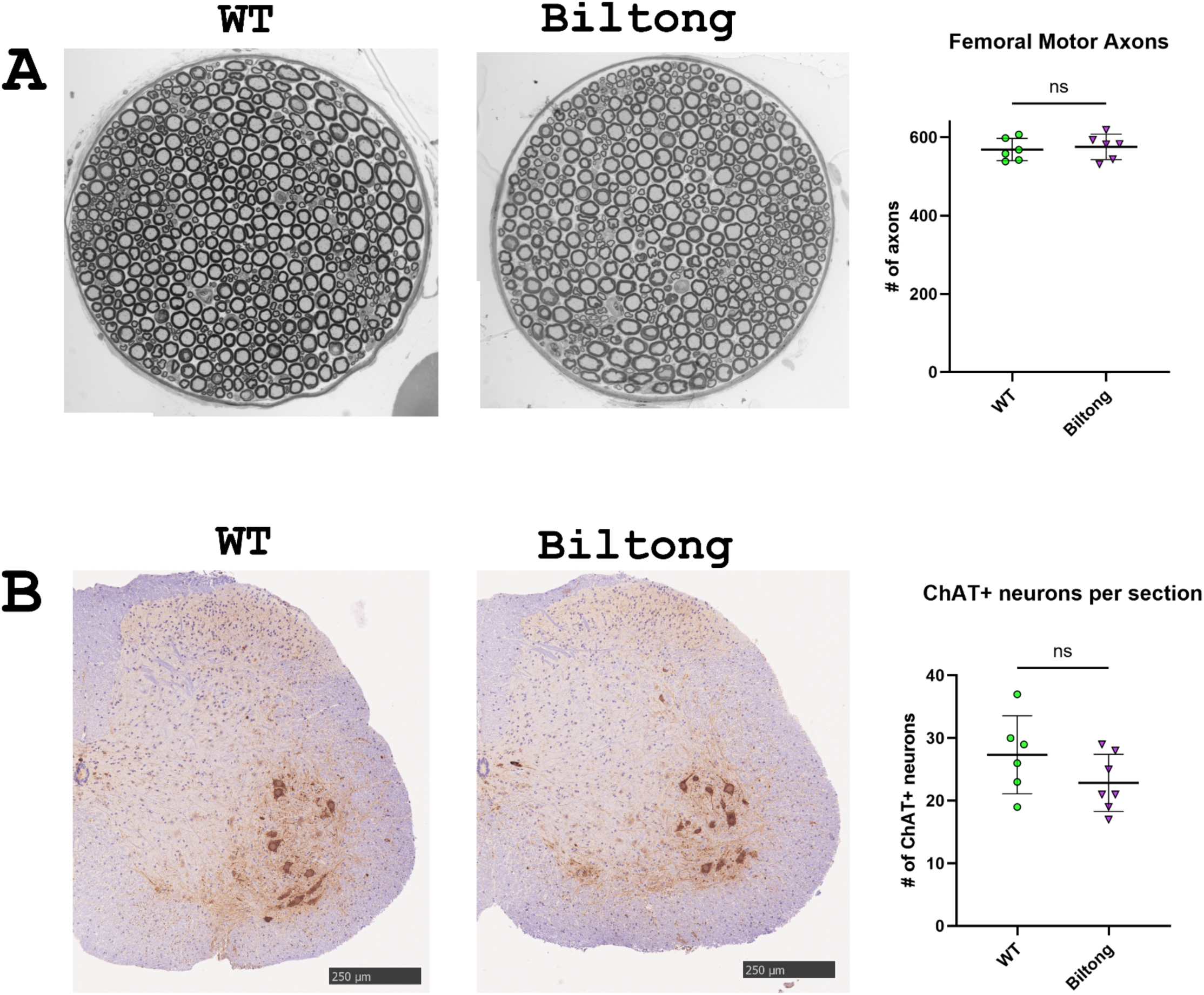
Biltong mice do not show signs of motor neuron degeneration at 6 months. **A)** Images of semithin toluidine blue-stained cross-sections of the motor branch of the femoral nerve and quantification of the number of axons in nerves from WT and Biltong mice at 6 months. N = 6 WT and 6 Biltong mice. **B)** Images of ChAT-labeled spinal cord cross-sections and quantification of motor neurons in spinal cord sections from WT and Biltong mice at 6 months. N = 6 WT and 7 Biltong mice.

**Supplementary Figure 3.**
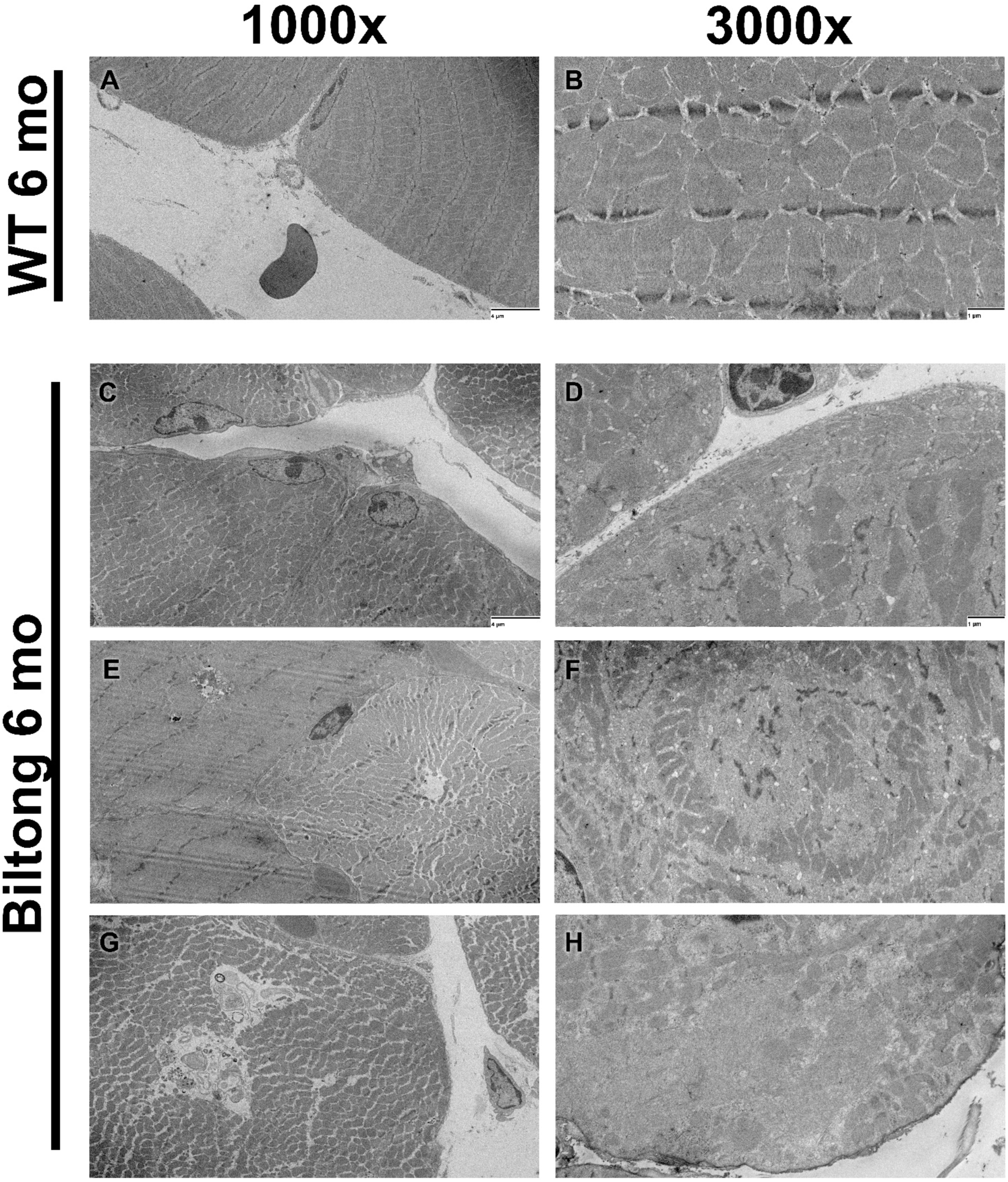
No true ringbinden seen with TEM on tibialis posterior from Biltong mice. **A-B)** Representative TEM images of tibialis posterior muscle from WT mice showing normal ultrastructure. **C-H)** TEM images of tibialis posterior muscle from Biltong mice revealed pathology similar to the soleus (Figure 6) including Z-band streaming, osmiophilic inclusions and multilamellar myeloid bodies, and numerous fibers with disorganized myofibrils. However, no fibers were seen with the typical perpendicular re-organization of myofibrils seen in true ringbinden observed in myotonic or limb-girdle muscular dystrophy patients.

**Supplementary Figure 4.**
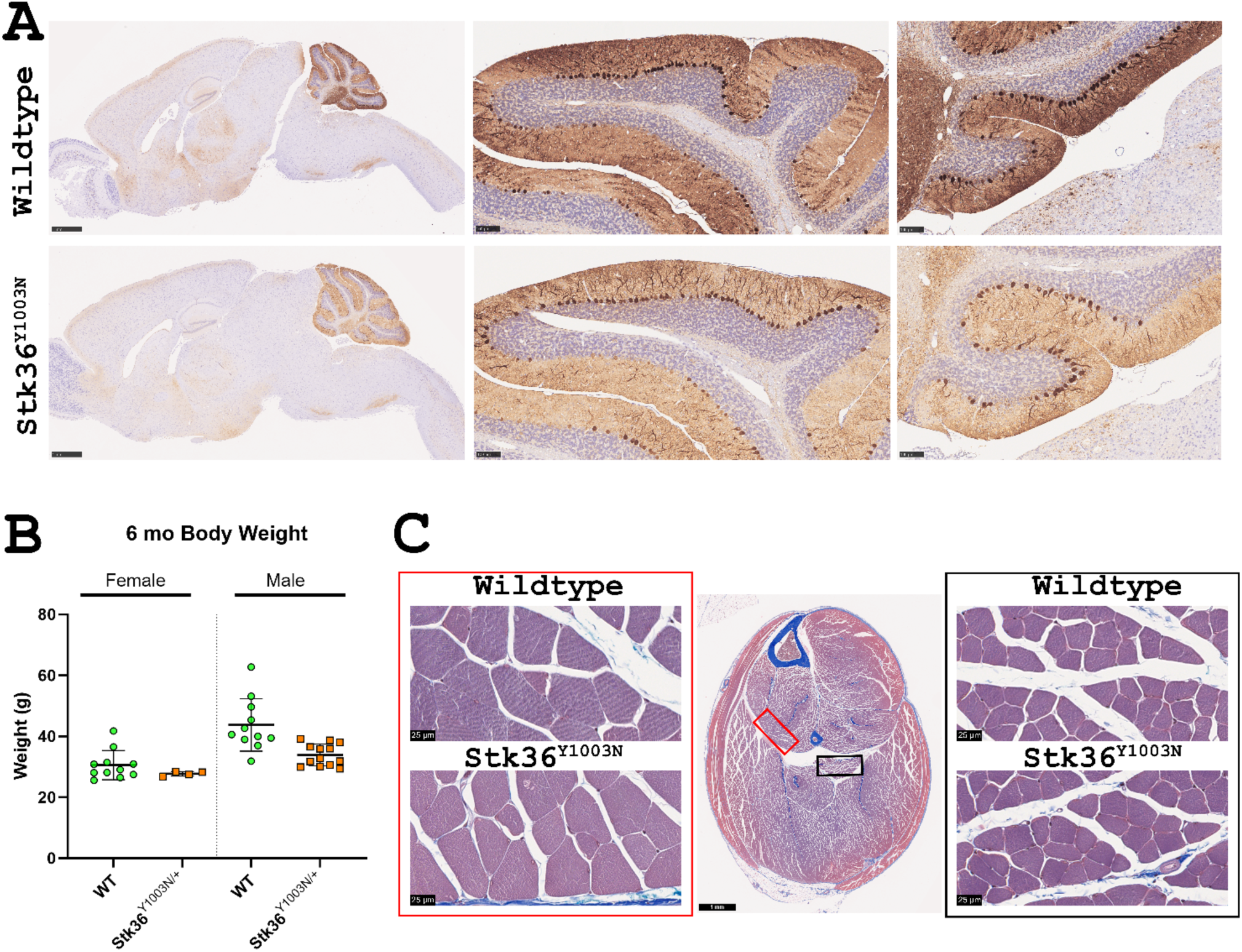
*Stk36^Y1003N^* knockin mouse does not produce a discernible phenotype by 6 months. **A)** Sagittal brain sections from 6 mo WT and *Stk36*^Y1003N^ knockin mice stained with Calbindin to label cerebellar Purkinje neurons. High magnification images show lobules VI (left) and X (right). Scale bars are 1 mm (whole brain) and 100 µm (lobules VI and X). **B)** Body weights from 6 mo WT and *Stk36*^Y1003N^ female and male mice. **C)** Masson’s trichrome staining of lower hindlimb cross-sections taken from WT and *Stk36*^Y1003N^ knockin mice at 6 months. High magnification (80x) insets are shown from the tibialis posterior (left, red rectangle) and soleus (right, black rectangle). Scale bars are 1 mm (whole hindlimb section) or 25 µm (80x images).

