## Supplemental Table 1 for "A mouse model of autosomal dominant spastic ataxia and myopathy caused by a mutation in *Tuba4a*"

| **Chromosome** | **Start** | **End** | **cM** | **strand GRCm39** | **MGI ID** | **Feature Type** | **Symbol** | **Name** |
| --- | --- | --- | --- | --- | --- | --- | --- | --- |
| 1 | 68071345 | 69147218 | 33.8 | - | MGI:104771 | protein coding gene | Erbb4 | erb-b2 receptor tyrosine kinase 4 |
| 1 | 69570373 | 69726404 | 34.9 | - | MGI:1342541 | protein coding gene | Ikzf2 | IKAROS family zinc finger 2 |
| 1 | 69866129 | 70764291 | 35.04 | + | MGI:1913972 | protein coding gene | Spag16 | sperm associated antigen 16 |
| 1 | 70764874 | 70924556 | 35.46 | + | MGI:2444069 | protein coding gene | Vwc2l | von Willebrand factor C domain-containing protein 2-like |
| 1 | 71066694 | 71142300 | 35.67 | - | MGI:1328361 | protein coding gene | Bard1 | BRCA1 associated RING domain 1 |
| 1 | 71282249 | 71454069 | 35.81 | - | MGI:2676312 | protein coding gene | Abca12 | ATP-binding cassette, sub-family A member 12 |
| 1 | 71596315 | 71618562 | 36.03 | + | MGI:1351352 | protein coding gene | Atic | 5-aminoimidazole-4-carboxamide ribonucleotide formyltransferase/IMP cyclohydrolase |
| 1 | 71624679 | 71692359 | 36.05 | - | MGI:95566 | protein coding gene | Fn1 | fibronectin 1 |
| 1 | 72198601 | 72251466 | 36.39 | - | MGI:2151839 | protein coding gene | Mreg | melanoregulin |
| 1 | 72298326 | 72323473 | 36.46 | - | MGI:2148199 | protein coding gene | Pecr | peroxisomal trans-2-enoyl-CoA reductase |
| 1 | 72323554 | 72342263 | 36.48 | + | MGI:2442781 | protein coding gene | Tmem169 | transmembrane protein 169 |
| 1 | 72346586 | 72434111 | 36.5 | + | MGI:104517 | protein coding gene | Xrcc5 | X-ray repair complementing defective repair in Chinese hamster cells 5 |
| 1 | 72466271 | 72576089 | 36.54 | - | MGI:2683550 | protein coding gene | Marchf4 | membrane associated ring-CH-type finger 4 |
| 1 | 72622410 | 72672293 | 36.72 | + | MGI:1859183 | protein coding gene | Smarcal1 | SNF2 related chromatin remodeling ATPase like 1 |
| 1 | 72682139 | 72739738 | 36.8 | - | MGI:2442559 | protein coding gene | Ankar | ankyrin and armadillo repeat containing |
| 1 | 72750449 | 72752972 | 36.87 | + | MGI:98068 | protein coding gene | Rpl37a | ribosomal protein L37a |
| 1 | 72863662 | 72891633 | 36.94 | + | MGI:96437 | protein coding gene | Igfbp2 | insulin-like growth factor binding protein 2 |
| 1 | 72897224 | 72914024 | 36.94 | - | MGI:96440 | protein coding gene | Igfbp5 | insulin-like growth factor binding protein 5 |
| 1 | 73054233 | 73055058 | 36.94 | - | MGI:98784 | protein coding gene | Tnp1 | transition protein 1 |
| 1 | 73949390 | 74163608 | 38.17 | - | MGI:104552 | protein coding gene | Tns1 | tensin 1 |
| 1 | 74164700 | 74187382 | 38.38 | + | MGI:3588214 | protein coding gene | Rufy4 | RUN and FYVE domain containing 4 |
| 1 | 74193153 | 74200405 | 38.41 | + | MGI:105303 | protein coding gene | Cxcr2 | C-X-C motif chemokine receptor 2 |
| 1 | 74230944 | 74233790 | 38.44 | - | MGI:2448715 | protein coding gene | Cxcr1 | C-X-C motif chemokine receptor 1 |
| 1 | 74275656 | 74307368 | 38.49 | + | MGI:1923959 | protein coding gene | Arpc2 | actin related protein 2/3 complex, subunit 2 |
| 1 | 74317709 | 74318783 | 38.53 | + | MGI:2653863 | protein coding gene | Gpbar1 | G protein-coupled bile acid receptor 1 |
| 1 | 74318999 | 74323897 | 38.53 | - | MGI:107809 | protein coding gene | Aamp | angio-associated migratory protein |
| 1 | 74324089 | 74392853 | 38.53 | + | MGI:1930773 | protein coding gene | Pnkd | paroxysmal nonkinesiogenic dyskinesia |
| 1 | 74327406 | 74343495 | 38.53 | - | MGI:1916910 | protein coding gene | Tmbim1 | transmembrane BAX inhibitor motif containing 1 |
| 1 | 74401272 | 74408482 | 38.54 | + | MGI:2685062 | protein coding gene | Catip | ciliogenesis associated TTC17 interacting protein |
| 1 | 74414354 | 74425221 | 38.54 | + | MGI:1345275 | protein coding gene | Slc11a1 | solute carrier family 11 (proton-coupled divalent metal ion transporters), member 1 |
| 1 | 74430668 | 74436444 | 38.54 | + | MGI:2654470 | protein coding gene | Ctdsp1 | CTD small phosphatase 1 |
| 1 | 74448543 | 74474719 | 38.54 | + | MGI:98930 | protein coding gene | Vil1 | villin 1 |
| 1 | 74474670 | 74583443 | 38.54 | - | MGI:2442483 | protein coding gene | Usp37 | ubiquitin specific peptidase 37 |
| 1 | 74545217 | 74570001 | 38.54 | + | MGI:1928902 | protein coding gene | Cnot9 | CCR4-NOT transcription complex, subunit 9 |
| 1 | 74581175 | 74605137 | 38.54 | + | MGI:107469 | protein coding gene | Plcd4 | phospholipase C, delta 4 |
| 1 | 74605490 | 74627308 | 38.54 | - | MGI:1924514 | protein coding gene | Zfp142 | zinc finger protein 142 |
| 1 | 74627448 | 74631602 | 38.54 | + | MGI:1914071 | protein coding gene | Bcs1l | BCS1 homolog, ubiquinol-cytochrome c reductase complex chaperone |
| 1 | 74632907 | 74640556 | 38.54 | - | MGI:1890215 | protein coding gene | Rnf25 | ring finger protein 25 |
| 1 | 74640604 | 74676053 | 38.54 | + | MGI:1920831 | protein coding gene | Stk36 | serine/threonine kinase 36 |
| 1 | 74700804 | 74740991 | 38.54 | + | MGI:1914784 | protein coding gene | Ttll4 | tubulin tyrosine ligase-like family, member 4 |
| 1 | 74752733 | 74777051 | 38.54 | + | MGI:88594 | protein coding gene | Cyp27a1 | cytochrome P450, family 27, subfamily a, polypeptide 1 |
| 1 | 74778081 | 74788380 | 38.54 | - | MGI:1891343 | protein coding gene | Prkag3 | protein kinase, AMP-activated, gamma 3 non-catalytic subunit |
| 1 | 74811051 | 74824481 | 38.54 | + | MGI:98960 | protein coding gene | Wnt6 | wingless-type MMTV integration site family, member 6 |
| 1 | 74831178 | 74843335 | 38.54 | + | MGI:108071 | protein coding gene | Wnt10a | wingless-type MMTV integration site family, member 10A |
| 1 | 74894188 | 74896891 | 38.54 | + | MGI:1330828 | protein coding gene | Cdk5r2 | cyclin dependent kinase 5, regulatory subunit 2 |
| 1 | 74920668 | 74924578 |  | - | MGI:2449712 | protein coding gene | Fev | FEV transcription factor, ETS family member |
| 1 | 74929093 | 74932302 | 38.54 | - | MGI:104336 | protein coding gene | Cryba2 | crystallin, beta A2 |
| 1 | 74941230 | 74974758 | 38.54 | - | MGI:2444274 | protein coding gene | Cfap65 | cilia and flagella associated protein 65 |
| 1 | 74984474 | 74990831 | 38.55 | - | MGI:96533 | protein coding gene | Ihh | Indian hedgehog |
| 1 | 75006505 | 75101870 | 38.56 | - | MGI:1922820 | protein coding gene | Nhej1 | non-homologous end joining factor 1 |
| 1 | 75102185 | 75110534 | 38.6 | - | MGI:104516 | protein coding gene | Slc23a3 | solute carrier family 23 (nucleobase transporters), member 3 |
| 1 | 75112406 | 75119374 | 38.6 | - | MGI:1916421 | protein coding gene | Cnppd1 | cyclin Pas1/PHO80 domain containing 1 |
| 1 | 75119432 | 75124553 | 38.61 | + | MGI:2388278 | protein coding gene | Retreg2 | reticulophagy regulator family member 2 |
| 1 | 75145290 | 75148270 | 38.61 | + | MGI:1916068 | protein coding gene | Zfand2b | zinc finger, AN1 type domain 2B |
| 1 | 75148361 | 75157036 | 38.62 | - | MGI:1921354 | protein coding gene | Abcb6 | ATP-binding cassette, sub-family B member 6 |
| 1 | 75157509 | 75168654 | 38.62 | - | MGI:2138446 | protein coding gene | Atg9a | autophagy related 9A |
| 1 | 75168795 | 75176031 | 38.62 | + | MGI:1098746 | protein coding gene | Ankzf1 | ankyrin repeat and zinc finger domain containing 1 |
| 1 | 75174880 | 75187457 | 38.63 | - | MGI:1921827 | protein coding gene | Glb1l | galactosidase, beta 1-like |
| 1 | 75187482 | 75192250 | 38.63 | + | MGI:1313271 | protein coding gene | Stk16 | serine/threonine kinase 16 |
| 1 | 75190872 | 75196509 | 38.63 | - | MGI:1095410 | protein coding gene | Tuba4a | tubulin, alpha 4A |
| 1 | 75196739 | 75208737 |  | - | MGI:2686470 | protein coding gene | A630095N17Rik | RIKEN cDNA A630095N17 gene |
| 1 | 75213050 | 75222336 | 38.64 | + | MGI:1928739 | protein coding gene | Dnajb2 | DnaJ heat shock protein family (Hsp40) member B2 |
| 1 | 75223671 | 75241146 | 38.64 | - | MGI:102765 | protein coding gene | Ptprn | protein tyrosine phosphatase receptor type N |
| 1 | 75248843 | 75255059 | 38.65 | - | MGI:1098222 | protein coding gene | Resp18 | regulated endocrine-specific protein 18 |
| 1 | 75285209 | 75294648 | 38.71 | - | MGI:1278328 | protein coding gene | Dnpep | aspartyl aminopeptidase |
| 1 | 75336973 | 75345223 | 38.85 | + | MGI:94885 | protein coding gene | Des | desmin |
| 1 | 75351941 | 75408964 | 38.88 | + | MGI:109282 | protein coding gene | Speg | SPEG complex locus |
| 1 | 75412574 | 75419823 | 39.04 | + | MGI:1916330 | protein coding gene | Gmppa | GDP-mannose pyrophosphorylase A |
| 1 | 75427080 | 75450987 | 39.08 | + | MGI:2652846 | protein coding gene | Asic4 | acid-sensing ion channel family member 4 |
| 1 | 75451213 | 75455951 | 39.14 | - | MGI:106576 | protein coding gene | Chpf | chondroitin polymerizing factor |
| 1 | 75456176 | 75462349 | 39.15 | + | MGI:2443133 | protein coding gene | Tmem198 | transmembrane protein 198 |
| 1 | 75462469 | 75483134 | 39.15 | - | MGI:2138628 | protein coding gene | Obsl1 | obscurin-like 1 |
| 1 | 75483872 | 75487010 | 39.16 | + | MGI:96569 | protein coding gene | Inha | inhibin alpha |
| 1 | 75498173 | 75513979 | 39.16 | + | MGI:1918978 | protein coding gene | Stk11ip | serine/threonine kinase 11 interacting protein |
| 1 | 75522688 | 75536075 | 39.16 | + | MGI:109350 | protein coding gene | Slc4a3 | solute carrier family 4 (anion exchanger), member 3 |
| 1 | 77343822 | 77491725 | 39.55 | - | MGI:98277 | protein coding gene | Epha4 | Eph receptor A4 |
| 1 | 78077904 | 78173771 | 39.79 | - | MGI:97487 | protein coding gene | Pax3 | paired box 3 |
| 1 | 78286982 | 78396926 | 40.23 | + | MGI:3589109 | protein coding gene | Sgpp2 | sphingosine-1-phosphate phosphatase 2 |
| 1 | 78394612 | 78465534 | 40.44 | - | MGI:1346035 | protein coding gene | Farsb | phenylalanyl-tRNA synthetase, beta subunit |
| 1 | 78473663 | 78488795 | 40.57 | - | MGI:2652858 | protein coding gene | BC035947 | cDNA sequence BC035947 |
| 1 | 78487628 | 78514810 | 40.59 | + | MGI:1915643 | protein coding gene | Mogat1 | monoacylglycerol O-acyltransferase 1 |
| 1 | 78635600 | 78645305 | 40.84 | + | MGI:2445092 | protein coding gene | Utp14b | UTP14B small subunit processome component |
| 1 | 78635600 | 78685462 | 40.84 | + | MGI:1921455 | protein coding gene | Acsl3 | acyl-CoA synthetase long-chain family member 3 |
| 1 | 78794628 | 78797749 | 40.89 | + | MGI:1891125 | protein coding gene | Kcne4 | potassium voltage-gated channel, Isk-related subfamily, gene 4 |
| 1 | 79412386 | 79417837 | 40.89 | - | MGI:103033 | protein coding gene | Scg2 | secretogranin II |
| 1 | 79584593 | 79649689 | 40.92 | - | MGI:1891304 | protein coding gene | Ap1s3 | adaptor-related protein complex AP-1, sigma 3 |
| 1 | 79679979 | 79753764 | 40.94 | - | MGI:1916618 | protein coding gene | Wdfy1 | WD repeat and FYVE domain containing 1 |
| 1 | 79753735 | 79759162 | 40.96 | + | MGI:1916413 | protein coding gene | Mrpl44 | mitochondrial ribosomal protein L44 |
| 1 | 79772038 | 79836382 | 40.97 | - | MGI:101780 | protein coding gene | Serpine2 | serine (or cysteine) peptidase inhibitor, clade E, member 2 |
| 1 | 80176416 | 80192050 | 41.19 | - | MGI:3026880 | protein coding gene | Fam124b | family with sequence similarity 124, member B |
| 1 | 80242640 | 80318197 | 41.24 | - | MGI:1347360 | protein coding gene | Cul3 | cullin 3 |
| 1 | 80427314 | 80439165 |  | - | MGI:5791097 | protein coding gene | Gm45261 | predicted gene 45261 |
| 1 | 80478790 | 80736244 | 41.41 | - | MGI:2146320 | protein coding gene | Dock10 | dedicator of cytokinesis 10 |
| 1 | 81054667 | 81319479 | 41.75 | + | MGI:2443135 | protein coding gene | Nyap2 | neuronal tyrosine-phophorylated phosphoinositide 3-kinase adaptor 2 |
| 1 | 82210822 | 82269137 | 42.0 | - | MGI:99454 | protein coding gene | Irs1 | insulin receptor substrate 1 |
| 1 | 82294178 | 82423087 | 42.08 | + | MGI:1924117 | protein coding gene | Rhbdd1 | rhomboid domain containing 1 |
| 1 | 82426144 | 82564570 | 42.2 | - | MGI:104687 | protein coding gene | Col4a4 | collagen, type IV, alpha 4 |
| 1 | 82564647 | 82699778 | 42.32 | + | MGI:104688 | protein coding gene | Col4a3 | collagen, type IV, alpha 3 |
| 1 | 82702611 | 82730115 | 42.45 | + | MGI:1922984 | protein coding gene | Mff | mitochondrial fission factor |
| 1 | 82734370 | 82746182 | 42.48 | - | MGI:1913511 | protein coding gene | Tm4sf20 | transmembrane 4 L six family member 20 |
| 1 | 82748545 | 82748964 |  | - | MGI:6096961 | protein coding gene | Gm47791 | predicted gene, 47791 |
| 1 | 82817204 | 82878903 | 42.55 | + | MGI:1333754 | protein coding gene | Agfg1 | ArfGAP with FG repeats 1 |
| 1 | 82891046 | 82891851 |  | + | MGI:1925172 | protein coding gene | A030005L19Rik | RIKEN cDNA A030005L19 gene |
| 1 | 82902638 | 82903433 |  | + | MGI:1925167 | protein coding gene | A030014E15Rik | RIKEN cDNA A030014E15 gene |
| 1 | 82908292 | 82909120 |  | - | MGI:3644079 | protein coding gene | Gm6217 | predicted gene 6217 |
| 1 | 82920280 | 82921109 |  | - | MGI:1925163 | protein coding gene | A030003K21Rik | RIKEN cDNA A030003K21 gene |
| 1 | 82932907 | 82933734 |  | - | MGI:3779750 | protein coding gene | Gm7544 | predicted gene 7544 |
| 1 | 82938007 | 82938645 |  | + | MGI:6097226 | protein coding gene | Gm47955 | predicted gene, 47955 |
| 1 | 82978269 | 82979201 |  | + | MGI:6097234 | protein coding gene | Gm47959 | predicted gene, 47959 |
| 1 | 82990244 | 83016169 | 42.65 | - | MGI:1931307 | protein coding gene | Slc19a3 | solute carrier family 19, member 3 |
| 1 | 83019245 | 83020201 |  | - | MGI:3779575 | protein coding gene | Krtap28-10 | keratin associated protein 28-10 |
| 1 | 83022653 | 83023667 |  | + | MGI:6097249 | protein coding gene | Gm47969 | predicted gene, 47969 |
| 1 | 83036007 | 83037099 |  | - | MGI:1925171 | protein coding gene | A030005K14Rik | RIKEN cDNA A030005K14 gene |
| 1 | 83038651 | 83039642 | 42.67 | + | MGI:1918636 | protein coding gene | Krtap28-13 | keratin associated protein 28-13 |
| 1 | 83094487 | 83096888 | 42.69 | + | MGI:1329031 | protein coding gene | Ccl20 | C-C motif chemokine ligand 20 |
| 1 | 83137473 | 83188295 | 42.7 | + | MGI:1923089 | protein coding gene | Daw1 | dynein assembly factor with WDR repeat domains 1 |
| 1 | 83233163 | 83385853 | 42.73 | - | MGI:1924879 | protein coding gene | Sphkap | SPHK1 interactor, AKAP domain containing |
| 1 | 84014014 | 84317550 | 43.14 | - | MGI:2138391 | protein coding gene | Pid1 | phosphotyrosine interaction domain containing 1 |
| 1 | 84347560 | 84673942 | 43.4 | - | MGI:2152889 | protein coding gene | Dner | delta/notch-like EGF repeat containing |
| 1 | 84667369 | 84674052 |  | + | MGI:7512541 | protein coding gene | Gm57795 | predicted gene, 57795 |
| 1 | 84698910 | 84818237 | 43.42 | - | MGI:1309481 | protein coding gene | Trip12 | thyroid hormone receptor interactor 12 |
| 1 | 84817562 | 84878208 | 43.43 | + | MGI:1289192 | protein coding gene | Fbxo36 | F-box protein 36 |
| 1 | 84883619 | 84912855 | 43.43 | - | MGI:1919031 | protein coding gene | Slc16a14 | solute carrier family 16 (monocarboxylic acid transporters), member 14 |
| 1 | 85065860 | 85088016 | 43.44 | - | MGI:3037746 | protein coding gene | Sp140l1 | Sp140 nuclear body protein like 1 |
| 1 | 85132922 | 85286736 |  | - | MGI:5621395 | protein coding gene | Gm38510 | predicted gene, 38510 |
| 1 | 85148753 | 85250268 |  | + | MGI:6721579 | protein coding gene | Gm53253 | predicted gene, 53253 |
| 1 | 85177334 | 85191507 | 43.45 | + | MGI:3644536 | protein coding gene | Gm7609 | predicted pseudogene 7609 |
| 1 | 85219007 | 85260602 | 43.44 | - | MGI:3612702 | protein coding gene | Sp140l2 | Sp140 nuclear body protein like 2 |
| 1 | 85257384 | 85267618 |  | + | MGI:6722185 | protein coding gene | Gm53567 | predicted gene, 53567 |
| 1 | 85411973 | 85412863 |  | - | MGI:6388834 | protein coding gene | Gm52955 | predicted gene, 52955 |
| 1 | 85477408 | 85478298 | 43.6 | + | MGI:3644077 | protein coding gene | Gm7592 | predicted gene 7592 |
| 1 | 85504620 | 85526538 | 43.6 | - | MGI:1923364 | protein coding gene | Sp110 | Sp110 nuclear body protein |
| 1 | 85528099 | 85572758 | 43.6 | + | MGI:3702467 | protein coding gene | Sp140 | Sp140 nuclear body protein |
| 1 | 85577709 | 85637719 | 43.6 | + | MGI:109561 | protein coding gene | Sp100 | nuclear antigen Sp100 |
| 1 | 85644804 | 85664377 | 43.75 | - | MGI:2443131 | protein coding gene | A630001G21Rik | RIKEN cDNA A630001G21 gene |
| 1 | 85721162 | 85779297 | 43.94 | + | MGI:107438 | protein coding gene | Cab39 | calcium binding protein 39 |
| 1 | 85822231 | 85836419 | 43.94 | + | MGI:1927594 | protein coding gene | Itm2c | integral membrane protein 2C |
| 1 | 85856204 | 85859477 | 43.94 | + | MGI:1918391 | protein coding gene | 4933407L21Rik | RIKEN cDNA 4933407L21 gene |
| 1 | 85866039 | 85888729 | 43.94 | - | MGI:2685064 | protein coding gene | Gpr55 | G protein-coupled receptor 55 |
| 1 | 85945728 | 85957683 | 43.94 | + | MGI:1917310 | protein coding gene | Spata3 | spermatogenesis associated 3 |
| 1 | 85973585 | 85983178 | 43.94 | + | MGI:1920042 | protein coding gene | 2810459M11Rik | RIKEN cDNA 2810459M11 gene |
| 1 | 85992341 | 86067017 | 43.94 | + | MGI:1917497 | protein coding gene | Psmd1 | proteasome (prosome, macropain) 26S subunit, non-ATPase, 1 |
| 1 | 86026748 | 86039692 | 43.94 | - | MGI:109323 | protein coding gene | Htr2b | 5-hydroxytryptamine (serotonin) receptor 2B |
| 1 | 86061359 | 86090510 |  | + | MGI:5439441 | protein coding gene | Gm21972 | predicted gene 21972 |
| 1 | 86082502 | 86206006 | 43.94 | + | MGI:1926045 | protein coding gene | Armc9 | armadillo repeat containing 9 |
| 1 | 86230943 | 86235027 | 43.94 | + | MGI:2384394 | protein coding gene | B3gnt7 | UDP-GlcNAc:betaGal beta-1,3-N-acetylglucosaminyltransferase 7 |
| 1 | 86272441 | 86287122 | 43.94 | - | MGI:97286 | protein coding gene | Ncl | nucleolin |
| 1 | 86313964 | 86317083 | 43.94 | - | MGI:1341898 | protein coding gene | Nmur1 | neuromedin U receptor 1 |
| 1 | 86354051 | 86355771 | 43.94 | + | MGI:1919113 | protein coding gene | Tex44 | testis expressed 44 |
| 1 | 86454448 | 86458434 | 43.96 | + | MGI:97803 | protein coding gene | Ptma | prothymosin alpha |
| 1 | 86470735 | 86510260 | 43.96 | - | MGI:1270843 | protein coding gene | Pde6d | phosphodiesterase 6D, cGMP-specific, rod, delta |
| 1 | 86510363 | 86534550 | 43.97 | + | MGI:1349388 | protein coding gene | Cops7b | COP9 signalosome subunit 7B |
| 1 | 86594015 | 86598295 | 43.98 | - | MGI:97369 | protein coding gene | Nppc | natriuretic peptide type C |
| 1 | 86631530 | 86977817 | 43.99 | + | MGI:2442555 | protein coding gene | Dis3l2 | DIS3 like 3'-5' exoribonuclease 2 |
| 1 | 87014416 | 87017650 | 44.05 | - | MGI:108009 | protein coding gene | Alppl2 | alkaline phosphatase, placental-like 2 |
| 1 | 87025724 | 87029328 | 44.05 | - | MGI:1924018 | protein coding gene | Alpi | alkaline phosphatase, intestinal |
| 1 | 87052695 | 87055634 | 44.06 | + | MGI:87984 | protein coding gene | Akp3 | alkaline phosphatase 3, intestine, not Mn requiring |
| 1 | 87075377 | 87084243 | 44.06 | - | MGI:1343461 | protein coding gene | Ecel1 | endothelin converting enzyme-like 1 |
| 1 | 87111035 | 87116127 | 44.07 | + | MGI:1916703 | protein coding gene | Prss56 | serine protease 56 |
| 1 | 87118329 | 87127792 | 44.07 | + | MGI:87893 | protein coding gene | Chrnd | cholinergic receptor, nicotinic, delta polypeptide |
| 1 | 87133533 | 87139365 | 44.07 | + | MGI:87895 | protein coding gene | Chrng | cholinergic receptor, nicotinic, gamma polypeptide |
| 1 | 87141636 | 87168210 | 44.08 | + | MGI:1914440 | protein coding gene | Eif4e2 | eukaryotic translation initiation factor 4E member 2 |
| 1 | 87192085 | 87238561 | 44.15 | + | MGI:1921607 | protein coding gene | Efhd1 | EF hand domain containing 1 |
| 1 | 87254720 | 87378518 | 44.23 | + | MGI:2138584 | protein coding gene | Gigyf2 | GRB10 interacting GYF protein 2 |
| 1 | 87314085 | 87322451 | 44.3 | - | MGI:3781032 | protein coding gene | Kcnj13 | potassium inwardly-rectifying channel, subfamily J, member 13 |
| 1 | 87397993 | 87403204 | 44.41 | + | MGI:1920484 | protein coding gene | Snorc | secondary ossification center associated regulator of chondrocyte maturation |
| 1 | 87404556 | 87501592 | 44.42 | - | MGI:1858414 | protein coding gene | Ngef | neuronal guanine nucleotide exchange factor |
| 1 | 87501749 | 87525567 | 44.44 | + | MGI:1344417 | protein coding gene | Neu2 | neuraminidase 2 |
| 1 | 87548034 | 87648229 | 44.44 | + | MGI:107357 | protein coding gene | Inpp5d | inositol polyphosphate-5-phosphatase D |
| 1 | 87683730 | 87720150 | 44.44 | + | MGI:1924290 | protein coding gene | Atg16l1 | autophagy related 16 like 1 |
| 1 | 87731402 | 87772880 | 44.44 | + | MGI:98227 | protein coding gene | Sag | S-antigen, retina and pineal gland (arrestin) |
| 1 | 87781009 | 87872902 | 44.44 | + | MGI:2138334 | protein coding gene | Dgkd | diacylglycerol kinase, delta |
| 1 | 87872841 | 87936273 | 44.45 | - | MGI:2443184 | protein coding gene | Usp40 | ubiquitin specific peptidase 40 |
| 1 | 87983110 | 88146726 | 44.49 | + | MGI:3580642 | protein coding gene | Ugt1a10 | UDP glycosyltransferase 1 family, polypeptide A10 |
| 1 | 87998522 | 88146719 | 44.5 | + | MGI:3576092 | protein coding gene | Ugt1a9 | UDP glucuronosyltransferase 1 family, polypeptide A9 |
| 1 | 88015550 | 88146720 | 44.5 | + | MGI:3576090 | protein coding gene | Ugt1a8 | UDP glucuronosyltransferase 1 family, polypeptide A8 |
| 1 | 88022784 | 88147724 | 44.5 | + | MGI:3032636 | protein coding gene | Ugt1a7c | UDP glucuronosyltransferase 1 family, polypeptide A7C |
| 1 | 88030979 | 88146720 | 44.51 | + | MGI:3580629 | protein coding gene | Ugt1a6b | UDP glucuronosyltransferase 1 family, polypeptide A6B |
| 1 | 88062531 | 88146719 | 44.52 | + | MGI:2137698 | protein coding gene | Ugt1a6a | UDP glucuronosyltransferase 1 family, polypeptide A6A |
| 1 | 88093734 | 88146719 | 44.53 | + | MGI:3032634 | protein coding gene | Ugt1a5 | UDP glucuronosyltransferase 1 family, polypeptide A5 |
| 1 | 88128323 | 88146719 | 44.54 | + | MGI:3576049 | protein coding gene | Ugt1a2 | UDP glucuronosyltransferase 1 family, polypeptide A2 |
| 1 | 88132454 | 88133470 | 44.55 | - | MGI:1306822 | protein coding gene | Dnajb3 | DnaJ heat shock protein family (Hsp40) member B3 |
| 1 | 88139681 | 88146719 | 44.55 | + | MGI:98898 | protein coding gene | Ugt1a1 | UDP glucuronosyltransferase 1 family, polypeptide A1 |
| 1 | 88154713 | 88190011 | 44.55 | + | MGI:3705228 | protein coding gene | Mroh2a | maestro heat-like repeat family member 2A |
| 1 | 88190193 | 88205355 | 44.57 | - | MGI:2685821 | protein coding gene | Hjurp | Holliday junction recognition protein |
| 1 | 88234457 | 88318909 | 44.57 | + | MGI:2181435 | protein coding gene | Trpm8 | transient receptor potential cation channel, subfamily M, member 8 |
| 1 | 88334683 | 88354160 | 44.63 | + | MGI:1922646 | protein coding gene | Spp2 | secreted phosphoprotein 2 |
| 1 | 88427593 | 88437788 | 44.67 | - | MGI:108038 | protein coding gene | Glrp1 | glutamine repeat protein 1 |
| 1 | 88625947 | 88629913 | 44.85 | - | MGI:2445172 | protein coding gene | Arl4c | ADP-ribosylation factor-like 4C |
| 1 | 88998137 | 89082790 | 45.05 | + | MGI:2138297 | protein coding gene | Sh3bp4 | SH3-domain binding protein 4 |
| 1 | 89382533 | 89823004 | 45.06 | + | MGI:2653690 | protein coding gene | Agap1 | ArfGAP with GTPase domain, ankyrin repeat and PH domain 1 |
| 1 | 89855684 | 89858898 | 45.06 | - | MGI:95668 | protein coding gene | Gbx2 | gastrulation brain homeobox 2 |
| 1 | 89880313 | 89942388 | 45.06 | - | MGI:2655109 | protein coding gene | Asb18 | ankyrin repeat and SOCS box-containing 18 |
| 1 | 89969854 | 90081123 | 45.08 | - | MGI:1922168 | protein coding gene | Iqca1 | IQ motif containing with AAA domain 1 |
| 1 | 90131702 | 90143446 | 45.28 | + | MGI:109562 | protein coding gene | Ackr3 | atypical chemokine receptor 3 |
| 1 | 90531147 | 90541063 | 45.53 | + | MGI:1915363 | protein coding gene | Cops8 | COP9 signalosome subunit 8 |
| 1 | 90694582 | 90771710 | 45.53 | - | MGI:88461 | protein coding gene | Col6a3 | collagen, type VI, alpha 3 |
| 1 | 90842807 | 90878864 | 45.73 | + | MGI:2176380 | protein coding gene | Mlph | melanophilin |
| 1 | 90880830 | 90881749 | 45.81 | + | MGI:3644668 | protein coding gene | Prlh | prolactin releasing hormone |
| 1 | 90885855 | 90897383 | 45.84 | - | MGI:104640 | protein coding gene | Rab17 | RAB17, member RAS oncogene family |
| 1 | 90926459 | 91056666 | 46.03 | + | MGI:1342770 | protein coding gene | Lrrfip1 | leucine rich repeat (in FLII) interacting protein 1 |
| 1 | 91072811 | 91098517 | 46.08 | + | MGI:2685663 | protein coding gene | Rbm44 | RNA binding motif protein 44 |
| 1 | 91107544 | 91152918 | 46.08 | + | MGI:1858418 | protein coding gene | Ramp1 | receptor (calcitonin) activity modifying protein 1 |
| 1 | 91177412 | 91214243 | 46.08 | + | MGI:1915171 | protein coding gene | Ube2f | ubiquitin-conjugating enzyme E2F (putative) |
| 1 | 91226060 | 91248797 | 46.08 | + | MGI:1355310 | protein coding gene | Scly | selenocysteine lyase |
| 1 | 91249797 | 91276028 | 46.08 | + | MGI:2685402 | protein coding gene | Espnl | espin-like |
| 1 | 91278795 | 91290126 | 46.09 | + | MGI:1918038 | protein coding gene | Klhl30 | kelch-like 30 |
| 1 | 91294152 | 91301939 | 46.1 | + | MGI:3606476 | protein coding gene | Erfe | erythroferrone |
| 1 | 91301583 | 91326537 | 46.11 | - | MGI:1914694 | protein coding gene | Ilkap | integrin-linked kinase-associated serine/threonine phosphatase 2C |
| 1 | 91339205 | 91341760 | 46.12 | - | MGI:1859852 | protein coding gene | Hes6 | hairy and enhancer of split 6 |
| 1 | 91343699 | 91387028 | 46.13 | - | MGI:1195265 | protein coding gene | Per2 | period circadian clock 2 |
| 1 | 91422369 | 91457029 | 46.16 | + | MGI:1921269 | protein coding gene | Traf3ip1 | TRAF3 interacting protein 1 |
| 1 | 91468266 | 91487311 | 46.19 | + | MGI:1929735 | protein coding gene | Asb1 | ankyrin repeat and SOCS box-containing 1 |
| 1 | 91729183 | 91775750 | 46.24 | + | MGI:104685 | protein coding gene | Twist2 | twist basic helix-loop-helix transcription factor 2 |
| 1 | 91856501 | 92123421 | 46.24 | - | MGI:3036234 | protein coding gene | Hdac4 | histone deacetylase 4 |
| 1 | 92367208 | 92401547 | 46.4 | - | MGI:1914523 | protein coding gene | Ndufa10 | NADH:ubiquinone oxidoreductase subunit A10 |
| 1 | 92407403 | 92408341 | 46.43 | - | MGI:3031250 | protein coding gene | Or6b2 | olfactory receptor family 6 subfamily B member 2 |
| 1 | 92418540 | 92419475 | 46.44 | - | MGI:3031249 | protein coding gene | Or6b2b | olfactory receptor family 6 subfamily B member 2B |
| 1 | 92438770 | 92446237 | 46.45 | - | MGI:3031248 | protein coding gene | Or6b3 | olfactory receptor family 6 subfamily B member 3 |
| 1 | 92500847 | 92501928 | 46.5 | + | MGI:3031247 | protein coding gene | Or9s23 | olfactory receptor family 9 subfamily S member 23 |
| 1 | 92516054 | 92517019 | 46.51 | + | MGI:3031246 | protein coding gene | Or9s27 | olfactory receptor family 9 subfamily S member 27 |
| 1 | 92524243 | 92525214 | 46.52 | + | MGI:3031245 | protein coding gene | Or9s15 | olfactory receptor family 9 subfamily S member 15 |
| 1 | 92535561 | 92536529 | 46.52 | + | MGI:3031244 | protein coding gene | Or9s14 | olfactory receptor family 9 subfamily S member 14 |
| 1 | 92547603 | 92548723 | 46.53 | + | MGI:107863 | protein coding gene | Or9s13 | olfactory receptor family 9 subfamily S member 13 |
| 1 | 92564867 | 92569707 | 46.55 | - | MGI:1914165 | protein coding gene | Cops9 | COP9 signalosome subunit 9 |
| 1 | 92571940 | 92576563 | 46.55 | - | MGI:2672814 | protein coding gene | Otos | otospiralin |
| 1 | 92759367 | 92787933 | 46.6 | + | MGI:1194891 | protein coding gene | Gpc1 | glypican 1 |
| 1 | 92787525 | 92830628 | 46.6 | - | MGI:3045261 | protein coding gene | Ankmy1 | ankyrin repeat and MYND domain containing 1 |
| 1 | 92834703 | 92836157 | 46.6 | + | MGI:1914696 | protein coding gene | Dusp28 | dual specificity phosphatase 28 |
| 1 | 92837697 | 92848307 | 46.6 | + | MGI:1914170 | protein coding gene | Rnpepl1 | arginyl aminopeptidase (aminopeptidase B)-like 1 |
| 1 | 92862130 | 92875670 | 46.6 | + | MGI:1344392 | protein coding gene | Capn10 | calpain 10 |
| 1 | 92878587 | 92914113 | 46.65 | + | MGI:1929509 | protein coding gene | Gpr35 | G protein-coupled receptor 35 |
| 1 | 92934056 | 92939991 | 46.72 | + | MGI:2664636 | protein coding gene | Aqp12 | aquaporin 12 |
| 1 | 92943186 | 93029673 | 46.74 | - | MGI:108391 | protein coding gene | Kif1a | kinesin family member 1A |
| 1 | 93062962 | 93073143 | 47.0 | + | MGI:1329033 | protein coding gene | Agxt | alanine-glyoxylate aminotransferase |
| 1 | 93079071 | 93088670 | 47.03 | - | MGI:1919124 | protein coding gene | Mab21l4 | mab-21-like 4 |
| 1 | 93096447 | 93158794 | 47.17 | + | MGI:3045962 | protein coding gene | Crocc2 | ciliary rootlet coiled-coil, rootletin family member 2 |
| 1 | 93163563 | 93228787 | 47.21 | + | MGI:3045960 | protein coding gene | Sned1 | sushi, nidogen and EGF-like domains 1 |
| 1 | 93228927 | 93233601 | 47.24 | - | MGI:1918355 | protein coding gene | Mterf4 | mitochondrial transcription termination factor 4 |
| 1 | 93237165 | 93271237 | 47.24 | - | MGI:2155936 | protein coding gene | Pask | PAS domain containing serine/threonine kinase |
| 1 | 93271350 | 93295344 | 47.24 | + | MGI:1913635 | protein coding gene | Ppp1r7 | protein phosphatase 1, regulatory subunit 7 |
| 1 | 93301652 | 93332025 | 47.24 | + | MGI:3052714 | protein coding gene | Ano7 | anoctamin 7 |
| 1 | 93333662 | 93406537 | 47.24 | - | MGI:99256 | protein coding gene | Hdlbp | high density lipoprotein (HDL) binding protein |
| 1 | 93406238 | 93437455 | 47.24 | + | MGI:97298 | protein coding gene | Septin2 | septin 2 |
| 1 | 93439826 | 93549698 | 47.24 | + | MGI:2385126 | protein coding gene | Farp2 | FERM, RhoGEF and pleckstrin domain protein 2 |
| 1 | 93548473 | 93581937 | 47.25 | - | MGI:1891699 | protein coding gene | Stk25 | serine/threonine kinase 25 (yeast) |
| 1 | 93613382 | 93623486 | 47.28 | + | MGI:1858494 | protein coding gene | Bok | BCL2-related ovarian killer |
| 1 | 93633113 | 93682560 | 47.29 | - | MGI:1914276 | protein coding gene | Thap4 | THAP domain containing 4 |
| 1 | 93682627 | 93717328 | 47.32 | + | MGI:1913865 | protein coding gene | Atg4b | autophagy related 4B, cysteine peptidase |
| 1 | 93720298 | 93729656 | 47.34 | - | MGI:108396 | protein coding gene | Dtymk | deoxythymidylate kinase |
| 1 | 93731687 | 93749823 | 47.34 | + | MGI:1922816 | protein coding gene | Ing5 | inhibitor of growth family, member 5 |
| 1 | 93752631 | 93780070 | 47.36 | + | MGI:2138209 | protein coding gene | D2hgdh | D-2-hydroxyglutarate dehydrogenase |
| 1 | 93789066 | 93804220 | 47.38 | + | MGI:2685834 | protein coding gene | Gal3st2 | galactose-3-O-sulfotransferase 2 |
| 1 | 93846159 | 93870367 | 47.41 | + | MGI:3711964 | protein coding gene | Gal3st2b | galactose-3-O-sulfotransferase 2B |
| 1 | 93918227 | 93939261 | 47.45 | + | MGI:3646771 | protein coding gene | Gal3st2c | galactose-3-O-sulfotransferase 2C |
| 1 | 93948215 | 93956056 | 47.46 | + | MGI:2661364 | protein coding gene | Neu4 | sialidase 4 |
| 1 | 93966027 | 93980278 | 47.47 | - | MGI:104879 | protein coding gene | Pdcd1 | programmed cell death 1 |
